# Phylogeny, classification, and character evolution of tribe Citharexyleae (Verbenaceae)

**DOI:** 10.1101/2020.10.08.331355

**Authors:** Laura A. Frost, Nataly O’Leary, Laura P. Lagomarsino, David C. Tank, Richard G. Olmstead

**Affiliations:** Department of Biology and Burke Museum, University of Washington, Seattle, WA, USA, 98195; Instituto de Botánica Darwinion, Labardén 200, San Isidro, Argentina; Department of Biological Sciences, Louisiana State University, Baton Rouge, LA, USA, 70803; Department of Biological Sciences, University of Idaho, Moscow, ID, USA, 83844

**Keywords:** *Baillonia*, *Citharexylum*, Citharexyleae, classification, microfluidic PCR, molecular phylogenetics, Neotropics, *Rehdera*, Verbenaceae

## Abstract

**Premise of the study:** Tribe Citharexyleae comprises three genera: *Baillonia, Citharexylum*, and *Rehdera*. While there is good support for these genera as a clade, relationships between genera remain unresolved due to low sampling of the largest genus, *Citharexylum*. A molecular phylogenetic approach was taken to resolve intergeneric relationships in Citharexyleae and infrageneric relationships in *Citharexylum*.

**Methods:** Seven chloroplast regions, two nuclear ribosomal spacers, and six low-copy nuclear loci were analyzed for 64 species of Citharexyleae. Phylogenetic analyses were conducted using maximum likelihood, Bayesian inference, and Bayesian multi-species coalescent approaches. Habit, presence/absence of thorns, inflorescence architecture, flower color, fruit color, and geography were examined to identify diagnostic characters for clades within *Citharexylum*.

**Key results:** Intergeneric relationships resolved *Rehdera* as sister to *Citharexylum* and *Baillonia* nested within *Citharexylum*. Two species, *C. oleinum* and *C. tetramerum*, fell outside of Citharexyleae close to tribe Duranteae. There is strong support for seven clades within *Citharexylum*, each characterized by a unique combination of geography, fruit color/maturation, and inflorescence architecture.

**Conclusions:** *Baillonia* is included in *Citharexylum; Rehdera* is retained as a distinct genus. A subgeneric classification for *Citharexylum* is proposed.

---

Prior to molecular phylogenetics, there was disagreement on the division and definition of groups within Verbenaceae. Authorities employing different criteria (e.g., gynoecial structure, inflorescence architecture, fruit characteristics) arrived at different divisions of the family (Atkins, 2004; Marx et al., 2010). Given the widespread conflict among taxonomists of the family, it is uncharacteristic that all morphology-based classifications of Verbenaceae recognized a group including *Citharexylum, Duranta*, and *Rhaphithamnus* (Bentham, 1839; Schauer, 1847; Briquet, 1895; Moldenke, 1971; Troncoso, 1974; Sanders, 2001; Atkins, 2004) on the basis of woody habit, fleshy fruit, and the presence of a staminode. Despite their strong morphological affinity, molecular studies indicate that each genus is more closely related to taxa that differ in one or more of these traits (Marx et al., 2010; Yuan et al., 2010; O’Leary et al. 2012).

Tribe Citharexyleae was recently recircumscribed to reflect evolutionary relationships (Marx et al., 2010). It now comprises *ca*. 133 spp. of Neotropical trees and shrubs in three genera: *Baillonia* (1 spp.), *Citharexylum* (130 spp.), and *Rehdera* (2 spp.). *Baillonia* is distributed in Brazil and Paraguay; *Citharexylum* is widespread throughout the Neotropics from Northern Mexico to Southern Brazil and Argentina, and *Rehdera* is distributed in Mesoamerica from the Yucatán peninsula in Mexico to the Nicoya peninsula in northwestern Costa Rica. All genera share a woody habit, extra-floral nectaries, terminal inflorescences, minute floral bracts, short pedicellate flowers, and the presence of a staminode. However, none of these traits are exclusive to these genera within Verbenaceae, and morphological synapomorphies that define the tribe have not been discovered (Marx et al. 2010; O’Leary et al., 2012).

The three genera of Citharexyleae are distinguished by carpel number and fruit type. *Citharexylum* and *Rehdera* have bi-carpellate ovaries that develop into fruits with two, two-seeded mericarps, while *Baillonia* has one carpel that aborts during development, giving rise to a uni-carpellate, two-seeded fruit (Briquet, 1895). *Baillonia* was traditionally classified in tribe Lantaneae, where all unicarpellate members of Verbenaceae were assigned before molecular phylogenies demonstrated the multiple origins of this trait. In addition to carpel number, these genera differ in whether their fruits are fleshy or dry: while *Citharexylum* and *Baillonia* both produce fleshy, drupaceous fruits, those of *Rehdera* are dry and dehiscent. Due to overall similarity in non-fruit traits, the dry-fruited species initially were classified as *Citharexylum* (Moore, 1895; Junell, 1934), but were segregated into *Rehdera* by Moldenke (1935).

Estimated to comprise 130 species (Atkins 2004), *Citharexylum* is by far the largest genus of Citharexyleae, as well as one of the largest genera of Verbenaceae. This would be true even if the suggestions that this estimate is inflated (Sanders, 2001; O’Leary and Frost, in review) are validated. In addition to the fleshy, bi-carpellate fruits that define the genus within the tribe, *Citharexylum* exhibits cryptic dioecy (Tomlinson and Fawcett, 1972; Rocca and Sazima, 2006; Rueda and Hammel, in prep.), with flowers that contain both stamens and pistils, but one reproductive whorl is sterile so that individuals are functionally staminate or pistillate. Dioecy is rare in Verbenaceae, known only in *Citharexylum* and *Lippia* section *Dioicolippia* (Múlgura, 2010). It is unknown whether dioecy is universal in *Citharexylum*. Despite the size of the genus, its taxonomy has been poorly studied and described. Moldenke published a series of notes (1957-1959) preparatory to a monograph that was never completed and Walpers (1845) proposed infrageneric groupings that were never validly published. These sections, *Spinosa* and *Inermia*, were defined based on presence or absence of spines, respectively; section *Inermia* was further divided by calyx morphology. Section *Spinosa* included five taxa that are recognized today as *C. flexuosum*, *C. montevidense*, and *Rhaphithamnus spinosus* (now distinct from *Citharexylum*); section *Inermia* encompassed the remaining 29 taxa in the treatment (Walpers, 1845). Moldenke (1957-1959) mirrored Walpers’ sections in a dichotomous key, updating the sections to include the many taxa described (mostly by Moldenke) since the informal treatment by Walpers. Moldenke placed *C. andinum, C. herrerae*, and *C. weberbaueri* in section *Spinosa*, leaving over 100 taxa—the vast majority of species in *Citharexylum—*listed in section *Inermia*. Due to the lack of structure within *Inermia*, this division provides no hypothesis of relationships between species, which vary in other traits including habit, flower color, inflorescence architecture, and fruit color.

Further phylogenetic study is necessary to fully understand the generic relationships within Citharexyleae and their implications for reclassification. The first phylogeny to include all three genera and sample multiple species of *Citharexylum* and *Rehdera*, supported inclusion of *Baillonia* within *Citharexylum* and a sister relationship of *Rehdera* to *Citharexylum* + *Baillonia* (Marx et al., 2010). However, because only 11 species in Citharexyleae were sampled and only chloroplast data were used, greater taxonomic sampling (especially of the large genus *Citharexylum*) and more loci are needed to fully resolve generic relationships. A more recent study expanded taxonomic sampling in *Citharexylum* and included one nuclear locus (Frost et al., 2017). This provided a framework phylogeny of *Citharexylum*, but did not test intergeneric relationships in the tribe.

We fill this knowledge gap by significantly expanding taxonomic and genetic sampling in a molecular phylogeny of Citharexyleae. Microfluidic PCR was used to target seven chloroplast regions, two nuclear rDNA spacers, and five low-copy nuclear genes. These data were analyzed in multiple frameworks, including a species-tree approach. With this improved sampling, we (1) resolve relationships between genera of Citharexyleae, (2) identify major clades of *Citharexylum* and the characters that unite them, (3) infer trait evolution within the tribe, and (4) establish a subgeneric classification for *Citharexylum*.

## MATERIALS AND METHODS

### Taxon sampling

One hundred eighty-two accessions representing 67 species of Citharexyleae were included in this study: one accession of monotypic *Baillonia*, 176 accessions of 64 *Citharexylum* species, and five accessions of two *Rehdera* species. Nine species from across Verbenaceae were included as outgroups. Vouchers and GenBank accession numbers may be found in Appendix S1 (see the Supplementary Data with this article). Herbarium acronyms follow Index Herbariorum (Thiers, constantly updated: http://sweetgum.nybg.org/science/ih/).

### Microfluidic PCR Primer Design and Validation

Targeted loci were selected for their phylogenetic utility and use in previous studies of Verbenaceae (O’Leary et al., 2009; Marx et al., 2010; Yuan et al., 2008, 2009a, 2009b; Lu-Irving and Olmstead, 2013; Lu-Irving et al. 2014; Frost et al., 2017). Seven chloroplast regions (cpDNA: *mat*K, *ndh*F, *rbc*L, *rpl*32, *rpo*C2, *trn*T-*trn*L, and *trn*L-*trn*F), two spacer regions (ETS and ITS) of nuclear 18S/26S rDNA (nrDNA), and five low-copy nuclear genes (waxy, PPR 24, PPR 62, PPR 70, and PPR 123) were included in this study.

Primer design and validation followed Uribe-Convers et al. (2016), with the exception that primer sequences were designed by eye, instead of using an R script, from alignments of published data (ncbi.nlm.nih.gov). Each primer pair was designed to amplify 400-600 bp. Validation reactions were performed with four experimental samples: two *Citharexylum* species (*C. schottii* and *C. myrianthum*) to test performance at the genus level, one taxon from the outgroup (*Duranta coriacea*) to test performance at the family level, and a negative control. Sixteen primer pairs were selected for seven chloroplast regions. One primer pair was used for each of the nrDNA and low-copy nuclear loci except for waxy, which was amplified with two primer pairs. Primer pair sequences can be found in Appendix S2 (see the Supplementary Data with this article).

### DNA extraction, Microfluidic PCR, and sequencing

Genomic DNA was extracted from field-collected, silica-gel-dried tissues using a modified CTAB method (Doyle and Doyle, 1987) or from herbarium specimens using a DNeasy plant mini kit (Qiagen, Valencia, California, USA). A total of 7603 bp of cpDNA were targeted for amplification and sequencing, including: 890 bp of *matK* in two segments, 1823 bp of *ndhF* in four segments, 1029 bp of *rbcL* in two segments, 525 bp of *rpl32* in one segment, 1893 bp of *rpoC2* in four segments, 512 bp of *trnT* in one segment, and 931 bp of *trnLF* in two segments. A total of 5132 bp of nuclear DNA were targeted for amplification and sequencing, including: 859 bp of nrDNA (400 bp of ETS and 459 bp of ITS) and 4274 bp for low-copy nuclear regions (332 bp of PPR24, 534 bp of PPR62, 341 bp of PPR70, 595 bp of PPR123, 967 bp of waxy haplotype 1, and 968 bp of waxy haplotype 2).

Microfluidic PCR was performed in two separate runs on an Access Array System (Fluidigm), one with a 48.48 Access Array integrated fluidic circuit (IFC) and another with a Juno 192.24 IFC. The same primer pairs were submitted for each IFC. Amplicons were harvested and pooled as described in Uribe-Convers et al. (2016). For each IFC, the resulting pools were multiplexed in an Illumina Miseq using the Reagent Kit v3 600 cycles. Microfluidic PCR in the Access Array System, downstream quality control, and Illumina sequencing were performed in the University of Idaho (Moscow, ID) Institute for Bioinformatics and Evolutionary Studies (ibest) Genomic Resources Core facility.

### Data processing

Reads from the Illumina Miseq runs were processed using the Fluidigm2PURC pipeline for processing paired-end Illumina data generated using the Fluidigm platform (Blischak et al., 2018). Reads were demultiplexed using the program dbcAmplicons (Uribe-Convers et al., 2016). Paired reads were separated by locus, with the fluidigm2purc script (github.com/pblischak/fluidigm2PURC). Each locus was processed using PURC’s purc_recluster.py script (github.com/pblischak/fluidigm2PURC; Edgar, 2004, 2010; Camacho et al., 2009; Edgar et al., 2011; Martin 2011; Rothfels et al., 2016) to iteratively run chimera detection and sequence clustering to produce a set of putative haplotypes. Reads were processed with clustering thresholds of 0.975, 0.99, 0.995, 0.997 and a size threshold of 5 (i.e. only clusters with ≥5 identical reads were retained). The clusters were run through the crunch-clusters script to infer haplotypes (github.com/pblischak/fluidigm2PURC). Chloroplast regions were processed as haploid. Reads from nuclear loci were processed with “unknown” ploidy, meaning all possible haplotypes were kept and haplotype number could vary by sample. Gene trees were generated with RAxML ver. 8.2.11 (Stamatakis, 2014) to process the haplotypes for nuclear loci. For individuals with haplotypes that nested together in gene trees (e.g., haplotypes that exhibited sequence similarity, but differed in amplicon length or a few heterogeneous sites) either the longest read or largest cluster was selected to represent the individual. For individuals with haplotypes that nested in different clades throughout the tree, all haplotypes were kept.

Raw reads for each locus were parsed into separate files by the purc_recluster.py script. In the instances in which raw data were available, but with insufficient coverage to meet the filters set in purc_recluster.py, the raw data were assessed and incorporated manually. A consensus sequence was generated from the available reads for regions with a single haplotype. For regions with multiple haplotypes, reads were analyzed individually in a phylogenetic context using RAxML ver. 8.2.11 (Stamatakis, 2014); consensus sequences were generated for reads that formed clades.

Four datasets were assembled for each region with different tiers of quality: 1) the original output from data processing, 2) the former plus individuals/haplotypes with ≥5 raw reads, 3) the former plus individuals/haplotypes with 2-4 raw reads, and 4) the former plus individuals/haplotypes with only a single read. Phylogenetic analyses were performed for each tier with RAxML ver. 8.2.11 (Stamatakis, 2014) to assess effects of the supplemented data on support values and topology. This supplementation technique was used only for individuals that had sufficient data to be included in the study, but had missing data for some loci due to low coverage. No additional individuals were added to the study as a result of these low-coverage reads.

### Phylogenetic analyses

#### Gene tree analyses

Sequences were aligned using AliView ver. 1.18.1 (Larsson, 2014); alignments were inspected and manually adjusted where necessary. The best-fit model of nucleotide substitution for each alignment was determined with jModelTest 2.1.7 (Darriba et al., 2012; Guindon and Gascuel, 2003); the GTR + G model was selected and applied to all alignments. Loci were concatenated using Geneious R7 7.0.6 (Biomatters, Auckland, New Zealand) and treated as a single dataset for regions with shared evolutionary histories: ndhF, rpoC2, trnT-trnL-trnF, rbcL, matK, and rpl32 are part of the haploid chloroplast genome (cpDNA), and ETS and ITS are tightly linked in the nuclear rDNA repeat (nrDNA). Each low-copy nuclear gene was treated as an independent locus. When multiple haplotypes were recovered for a region, haplotypes were separated and treated as independent loci. Eight independent loci were analyzed: cp, nr, PPR24, PPR62, PPR70, PPR123, waxy haplotype1, and waxy haplotype 2.

Phylogenetic analyses were conducted for individual datasets, low-copy nuclear genes combined, and all data combined using maximum likelihood and Bayesian approaches, as implemented in RAxML and MrBayes ver. 3.2.6 (Ronquist & Huelsenbeck, 2003), respectively. Maximum likelihood analyses consisted of bootstrap analyses with 500 replicates for each dataset. Datasets were partitioned into individual gene regions with individual parameters unlinked in Bayesian analyses. Analyses consisted of two replicate runs, each with four chains, which were run for at least one million generations, sampled every 200 generations. Runs were assessed for convergence and stationarity in Tracer v1.6.0 (Rambaut et al., 2014); our benchmark for assessing convergence were effective sample sizes (ESS) above 200 for all parameter values across combined files. The first 20% of sampled trees were discarded as burnin from each run; the remaining trees were summarized into a majority rule consensus tree with all compatible groups.

#### Species tree analyses

Species tree estimation was performed on the ingroup using the multispecies coalescent (MSC) model implemented in *BEAST (BEAST v1.8.4; Drummond et al. 2012). Substitution models and clock models were unlinked for the fourteen individual alignments. The GTR+G substitution model, empirical base frequencies, and a random local clock (Drummond and Suchard, 2010) were applied to each. Trees were linked based on shared evolutionary history, in the case of cp and nr datasets, or based on gene tree topology for low-copy nuclear genes. Two partitions were set for low-copy nuclear genes: (1) those where Mesoamerican species formed a clade (PPR24, PPR123, waxy haplotype 2) and (2) those where Mesoamerican species were paraphyletic with respect to South American species (PPR62, PPR70, waxy haplotype 1). Chloroplast data were treated as an organellar (haploid) locus with half the effective population size of a bi-parentally inherited locus, and the nuclear loci were treated as autosomal nuclear. The birth-death process was used as the species tree prior. Two replicate runs were performed for 500 million generations, sampling every 25,000. Convergence between runs and stationarity within were assessed in Tracer v1.6.0 (Rambaut et al., 2014); the duplicate runs were considered to have converged when effective sample sizes (ESS) for parameter values were above 200 for combined files. The first 10% of sampled tree states were discarded from each run as burn-in. Accessions of *C. bourgeauianum*, *C. flabellifolium*, *C. guatemalense*, and *C. reticulatum* were excluded from the species tree analyses due a high proportion of missing data. Additionally, *Citharexylum oleinum* and *C. tetramerum* were excluded because they were inferred to belong outside of the ingroup.

#### Morphology

In order to understand character evolution and describe clades in *Citharexylum*, morphological characters were examined. Targeted characters included habit, presence/absence of thorns, inflorescence architecture, flower number, corolla color, fruit color, and geography. These traits, which are easily observable in the field, from photographs, and from herbarium specimens, were chosen because they are practical, likely informative in recognizing large clades, and have the potential to fill gaps in natural history knowledge. For habit, species were scored as shrubs (i.e., less than six m tall and with several main stems) or trees (i.e., a single trunk and/or taller than 6 m). To best describe variation in inflorescence architecture terminal and axillary inflorescences were scored separately; these were scored as absent, simple, or compound. Average flower number per inflorescence was categorized as >20, 10-20, or <10. Corolla color was categorized as solid white, solid yellow, solid purple, or white/yellow with purple/orange on the abaxial surface of the petal lobes. Fruit color/maturation was categorized as red synchronous (green to red), black synchronous (green to black), black sequential (green to red to black) in *Citharexylum*. Because fruit type does not change within *Citharexylum* and fruit color and mode of maturation in *Citharexylum* was the focus of this analysis, the dry fruits of sister genus *Rehdera* were coded as brown. Geographic range was divided into three regions: Mesoamerica, Caribbean, and South America. Character data were pooled from Moldenke (1957-1959); field observations; herbarium records via personal visits to A/GH, F, MEXU, MO, and USM; and online databases GBIF (https://www.gbif.org/), CV Starr Virtual Herbarium (NYBG; sweetgum.nybg.org/science/vh/), SEINET (http://swbiodiversity.org/seinet/), and Tropicos (MOBOT; https://www.tropicos.org/).

#### Ancestral State Reconstructions

Ancestral character reconstructions were performed for all traits except geography using parsimony and maximum likelihood estimation. Parsimony reconstructions were performed using the “Trace Character History” > “Parsimony Ancestral States” function in Mesquite v3.51 (Maddison and Maddison, 2018). Maximum likelihood estimations were conducted using the R package phytools (Revell, 2012) using the ace function and an equal rates (ER) model. The biogeography of *Citharexylum* will be addressed in a separate, more detailed study using more appropriate methods for biogeographic reconstruction. Because the presence and/or complexity of terminal and/or axillary inflorescences varied within species, each inflorescence type was simply scored as present or absent for ancestral character reconstructions.

#### Tests for correlated evolution

The BayesDiscrete module in BayesTraits (Pagel and Meade, 2006) was used to test for correlation between morphological characters. Maximum likelihood analyses were conducted for each possible pair of characters coded as binary variables. Traits that were coded as multi-state in ancestral state reconstructions—flower number, flower color and fruit color/maturation—were simplified to binary variables or excluded. Flower number was scored as ≤20 flowers per inflorescence vs. ≥20 flowers per inflorescence. Flower color was scored pale or purple, including species that have purple lobes. Fruit maturation was simplified to synchronous or sequential. Fruit color was not included in correlation tests due to the difficulty of scoring red, black, and red-to-black fruits in a binary and biologically accurate way.

For each pair of characters, two maximum likelihood models were fit to the phylogeny in BayesTraits: an independent model (I) and a dependent model (D). The independent model assessed the likelihood of the data under the null hypothesis that the traits evolved independently, whereas the dependent model assumed the transition rates for one character were dependent upon the state of the other character. To determine if traits were correlated, the likelihood estimate of the independent model was compared to the likelihood estimate of the dependent model via a likelihood ratio test. The likelihood ratio was tested against a χ^2^ distribution, with four degrees of freedom (following Pagel 1994). Probabilities were calculated from χ^2^ values.

## RESULTS

### Dataset characteristics

The total concatenated dataset consisted of 12927 aligned bp and 176 accessions, including 167 accessions in Citharexyleae. Among the characters, 4140 were variable, of which 2398 were potentially parsimony-informative. One hundred thirty-nine accessions had sequence data for at least five of the eight loci, with 82 of these having data for at least seven loci. All accessions included in the concatenated dataset had data for both cpDNA and nrDNA. Nine accessions of *Citharexylum* with insufficient data to be included in the concatenated analyses, but representing species not otherwise included, were retained in individual gene tree analyses. Characteristics of individual loci are shown in Table 1.

**Table 1.** Characteristics of individual datasets

| Locus | No. of taxa | Proportion of taxa | Seq. length range (bp) | Alignment length (bp) | % missing data | No. of variable sites | Proportion of variable sites | No. of parsimony informative sites | Proportion of parsimony informative sites | GC content |
| --- | --- | --- | --- | --- | --- | --- | --- | --- | --- | --- |
| cpDNA | 179 | 0.923 | 1,623-7,575 | 7788 | 18.367 | 1782 | 0.229 | 922 | 0.118 | 0.362 |
| nrDNA | 188 | 0.969 | 393-859 | 865 | 9.946 | 591 | 0.683 | 499 | 0.577 | 0.621 |
| PPR 24 | 143 | 0.737 | 329-331 | 332 | 0.607 | 158 | 0.476 | 125 | 0.377 | 0.413 |
| PPR 62 | 106 | 0.546 | 358-534 | 534 | 4.111 | 208 | 0.39 | 109 | 0.204 | 0.406 |
| PPR 70 | 90 | 0.464 | 338-341 | 341 | 0.065 | 158 | 0.463 | 93 | 0.273 | 0.382 |
| PPR 123 | 82 | 0.423 | 375-536 | 595 | 12.494 | 244 | 0.41 | 113 | 0.19 | 0.398 |
| waxy hap 1 | 151 | 0.778 | 359-842 | 967 | 28.171 | 413 | 0.427 | 256 | 0.265 | 0.384 |
| waxy hap 2 | 140 | 0.722 | 388-906 | 968 | 37.093 | 462 | 0.477 | 297 | 0.307 | 0.381 |

### Phylogenetic analyses

#### Gene tree analyses

All gene trees support the monophyly of Citharexyleae, except for two species (*C. oleinum* and *C. tetramerum*), which nest among the outgroups. As in previous studies (Marx et al, 2010), *Rehdera* is inferred to be sister to *Citharexylum*, while *Baillonia* is nested within *Citharexylum* in all trees. *Citharexylum altamiranum* (clade C) is sister to the rest of the genus (Figs. 2 and 3), which is comprised of two large clades, A and B (Fig. 3). Six subclades within A and B are consistently found in individual and combined gene trees, but relationships among these varied. Terminal clades within *Citharexylum* with high support for all datasets included: (I) a large clade including *C. ellipticum*, *C. hidalgense*, *C. moccinoi*, *C. roxanae*, and *C. schottii*; (II) a clade composed of *C. brachyanthum*, *C. lycioides*, *C. racemosum* and *C. rosei*; (III) a clade composed of *C. affine*, and *C. hintonii*; (IV) a clade including *Baillonia amabilis, C. microphyllum*, and *C. montanum*; (V) a clade composed of *C. alainii* and *C. microphyllum*; and (VI) a clade/grade including *C. dentatum*, *C. flexuosum*, *C. herrerae*, and *C. montevidense*; and (VII = clade C) *C. altamiranum* (Figs. 2 and 3, but see Fig. 3 for tip labels). Clade A includes clades I, II, and III in the chloroplast, low-copy nuclear combined, and all data combined trees; the placement of clades II and III varied in individual gene trees. In all trees, clade B included clades IV, V, and VI (species in the latter often form a grade in individual gene trees), though relationships among taxa in clade VI differ among datasets (Appendices S3-S12). Due to the geographic distribution of most of their species, clade A (including clades I II, and III, as supported by combined analyses) will be referred to as the Mesoamerican clade and clade B as the South American clade.

*Citharexylum spinosum*, the type species for the genus, nested within different clades for different datasets. All three accessions nested within the Mesoamerican clade in cpDNA, PPR70, and PPR123 trees. Accessions were split, with one or more nesting in the Mesoamerican clade and one or more in the South American clade in nrDNA, PPR24, and PPR62 trees. Two accessions of *C. spinosum* had an additional copy for each of the waxy of haplotypes (i.e. four total copies of the waxy gene, two copies of haplotype 1 and two copies of haplotype 2). For each haplotype, one copy nested within the Mesoamerican clade and another nested within the South American clade (Appendices S9, S10). One accession also showed this pattern in PPR62 (*C. spinosum*_JCS5727; Appendix S6).

#### Species tree analyses

The species tree topology was consistent with the topology of the all-combined dataset (Fig. 4). The split between the Mesoamerican clade and the South American clade was supported in the species tree. The same seven clades were inferred in the species tree as in the individual and combined gene trees with high support. Relationships within subclade VI varied between replicate runs in the multispecies coalescent and differed from relationships recovered by the all combined dataset. Three clades within clade VI were consistently inferred in the species tree and all combined analyses: a clade of *C. flexuosum, C. kobuskianum, C. peruvianum*, and *C. weberbaueri*, a clade of *C. andinum, C. joergensenii*, and *C. montevidense*, and a clade including *C. argutedentatum, C. herrerae*, and *C. sulcatum*.

#### Morphology

Observations were available for all characters for all species except fruit color in *C. racemosum* (Fig. 4). Variation within species was observed for traits related to inflorescence architecture. Terminal inflorescences were present in all species, but often absent in individuals of *C. subflavescens* and *C. montanum*. Terminal inflorescence complexity (i.e., simple or compound) varied within all species of *Citharexylum* for which compound terminal inflorescences were observed; only *Rehdera penninervia* was found to consistently have compound terminal inflorescences. Similarly, there was variation within species for presence/absence of axillary inflorescences. Individuals of 27 of the species—nearly half the taxonomic diversity of the tribe—had axillary inflorescences. Of those 27, 12 species were observed to consistently have axillary inflorescences, and six included individuals with compound axillary inflorescences. As with the terminal infloresence, *Rehdera penninervia* was the only species for which members consistently had compound axillary inflorescences.

#### Ancestral State Reconstructions

Habit type shifted repeatedly across the phylogeny; and proved difficult to reconstruct at deeper nodes (Appendix S13). Terminal inflorescences, which were present in all species but often absent in *C. montanum* and *C. subflavescens*, either became labile in the most recent common ancestor of *C. montanum, C. subflavescens*, and *C. rimbachii* and that lability was lost in C. rimbachii or lability was independently gained in the two species (Appendix S14). Axillary inflorescences were lost in the ancestor of the two major clades of *Citharexylum* and regained independently nine times (Appendix S15). Reduction of flower number per inflorescence from >20 to <20 was estimated to have occurred in *C. altamiranum*, *C. alanii* and *C. microphyllum*, clade II, and clade VI (Appendix S16). Flowers with at least partial purple coloration arose six times throughout the evolutionary history genus (Appendix S17). Reconstructions of the evolutionary history of fruit color were inconsistent. Red synchronous fruits were estimated to be the ancestral fruit color in *Citharexylum*, with black synchronous fruits evolving once in *C. altamiranum* and one to three times in clade VI of the South American clade by maximum likelihood analyses (Fig. 5, Appendix S18A). Maximum parsimony reconstructions, however, suggested that black synchronous fruits were the ancestral state for the genus and the South American clade, with red synchronous fruits arising three times in clade VI (Appendix S18B). Likewise, maximum likelihood reconstructions had higher support for multiple origins of black sequential fruits in the Mesoamerican clade, whereas maximum parsimony reconstructions supported a single origin and multiple losses (Appendix S18).

The variable topologies of subclade VI have implications for character evolution of thorns (Appendix S19). Thorns are exclusively found in members of subclade VI, though not all species bear thorns. There was variable support for a single evolution of thorns in the ancestor of subclade VI followed by multiple losses (sometimes with secondary regains), or multiple independence evolutions of thorns (Appendix S20). Support for these reconstructions approached equivocal.

#### Tests for correlated evolution

Significant support was found for correlated evolution (i.e., dependent model favored over independent at p < 0.05) for five pairs of traits (Table 2): axillary inflorescence and flower number (p=0.0011); axillary inflorescence and fruit maturation (p=0.0064); axillary inflorescence and habit (p=0.0002); axillary inflorescence and thorns (p=0.0379); and habit and flower number (p=0.0020).

**Table 2.** Results from tests for correlated evolution. Significant p values (p < 0.05) and their corresponding characters are in bold.

| Characters | L(D) | L(I) | $\chi^2$ | p value |
| --- | --- | --- | --- | --- |
| axillary inflorescence, flower color | -53.4615 | -56.2591 | 5.5951 | 0.2315 |
| **axillary inflorescence, flower number** | -40.1691 | -49.3196 | 18.3010 | **0.0011** |
| **axillary inflorescence, fruit maturation** | -60.5637 | -67.7133 | 14.2992 | **0.0064** |
| **axillary inflorescence, habit** | -57.1479 | -68.1717 | 22.0478 | **0.0002** |
| axillary inflorescence, terminal inflorescence | -39.9981 | -41.8457 | 3.6952 | 0.4488 |
| **thorns, axillary inflorescence** | -41.6941 | -46.7707 | 10.1533 | **0.0379** |
| flower color, fruit maturation | -52.9804 | -54.6319 | 3.3030 | 0.5085 |
| flower number, flower color | -35.4795 | -36.2382 | 1.5174 | 0.8236 |
| flower number, fruit maturation | -45.5076 | -47.6924 | 4.3696 | 0.3583 |
| habit, flower color | -52.1010* | -55.0903 | 5.9786 | 0.2008 |
| **habit, flower number** | -39.6639* | -48.1508 | 16.9739 | **0.0020** |
| habit, fruit maturation | -63.8734* | -66.5445 | 5.3422 | 0.2540 |
| habit, terminal inflorescence | -38.7818* | -40.6769 | 3.7902 | 0.4351 |
| habit, thorns | -43.6396* | -45.6019 | 3.9247 | 0.4163 |
| terminal inflorescence, flower color | -28.2228* | -28.7643 | 1.0830 | 0.8970 |
| terminal inflorescence, flower number | -20.8600* | -21.8248 | 1.9297 | 0.7487 |
| terminal inflorescence, fruit maturation | -39.5042* | -40.2185 | 1.4286 | 0.8392 |
| thorns, flower color | -32.8826* | -33.6893 | 1.6134 | 0.8064 |
| thorns, flower number | -24.0216* | -26.7498 | 5.4565 | 0.2436 |
| thorns, fruit maturation | -43.2774* | -45.1435 | 3.7321 | 0.4435 |
| thorns, terminal inflorescence | -18.8144* | -19.2759 | 0.9229 | 0.9212 |

## CLASSIFICATION

Citharexyleae Briq. in H.G.A. Engler & K.A.E. Prantl, Nat. Pflanzenfam. IV, 3a: 144, 158. 1895. I. *Citharexylum* L., Sp. Pl. 1: 625. 1753. Type: *Citharexylum spinosum* L.

1. C. subgenus ***Citharexylum*** Type: *Citharexylum spinosum* L. section ***Citharexylum*** Trees and shrubs, thorns absent; inflorescences terminal or terminal and axillary, simple or compound; flowers many, corolla white, pale yellow-green, or purple; fruits maturing from orange to black. Distribution: Most species in this clade are distributed only in Mesoamerica; however, *C. spinosum* and *C. caudatum* extend into the Caribbean and South America. Several other species belonging to this clade—*C. karstenanii*, *C. subthyrsoideum*, and *C. decorum*— occur only in South America. Estimated species: 25-30 section ***Mexicanum*** L.A. Frost, sect. nov. Type: *Citharexylum brachyanthum* (A.Gray ex Hemsl.) A.Gray Shrubs, thorns absent; inflorescences terminal and simple, terminating very short branchlets; flowers few, corolla white; fruits red-orange at maturity or maturing from orange to black. Distribution: Members of this clade are mostly endemic to Mexico; they either border or occur within the Chihuahuan desert. *Citharexylum lycioides*, *C. racemosum*, and *C. rosei* are distributed along the southern/south-eastern edge of the Chihuahuan desert, whereas *C. brachyanthum* occurs throughout, rarely extending into Texas in the United States. Estimated species: 4 section ***Pluriflorum*** L.A. Frost, sect. nov. Type: *Citharexylum affine* D. Don. Trees, thorns absent; inflorescences terminal and axillary, simple or compound; flowers many, corolla purple; fruits maturing from orange to black. Distribution: Whereas *C. hintonii* is narrowly endemic to Mexico—most collections coming from the southwestern portion of the state of México—*C. affine* is more widespread throughout Mesoamerica. The latter species occurs throughout southern Mexico, Guatemala, Belize, and potentially Nicaragua (based on a few records from Nicaragua, but no known specimens from El Salvador or Honduras). Estimated species: 2-3
2. C. subgenus ***Sudamericanum*** L.A. Frost, subg. nov. Type: *C. flexuosum* (Ruiz & Pav.) D. Don. section ***Sylvaticum*** L.A. Frost, sect. nov. Type: *Citharexylum myrianthum* Cham. Trees or shrubs, thorns absent; inflorescences terminal, axillary, or terminal and axillary, simple or compound; flowers many, corolla white; fruits red. Distribution: Members of this clade are distributed widely throughout South America. Species typically occur in mesic forests and wet areas, from mid-elevation cloud forests in the northern Andes to moist lowlands surrounding the Amazon as well as those in southeastern Brazil, Paraguay, and Argentina. *Citharexylum amabilis* is distributed in the cerrado of Brazil; it is the only species of *Citharexylum* to occur in the cerrado. Estimated species: 12 – 15 section ***Caribe*** L.A. Frost, sect. nov. Type: *Citharexylum microphyllum* E.O. Schulz Shrubs, thorns absent; inflorescences terminal, simple; flowers few, corolla white; fruits red. Distribution: These species are endemic to the Greater Antilles; they are often found on limestone outcroppings, rocky slopes and hillsides. Estimated species: 2-7 section ***Armatum*** L.A. Frost, subg. nov. Type: Type: *C. flexuosum* (Ruiz & Pav.) D. Don. Shrubs, thorns present or absent; inflorescences terminal and simple; flowers few to submany, corollas white to pale yellow-green, some with lobes tipped purple or orange in bud; fruits red or black. Distribution: Members of this clade are largely distributed in the central Andes in Peru, Bolivia, and Argentina. *Citharexylum sulcatum* and *C. illicifolium* are distributed in Colombia and Ecuador, respectively. Species occur in arid to semi-arid inter-Andean valleys and high-elevation Andean grasslands. Estimated species: 14-16
3. C. subgenus ***Purpuratum*** L.A. Frost, subg. nov. Type: *Citharexylum altamiranum* Greenm. Shrub, thorns absent; inflorescences terminal and axillary, simple; flowers few, corollas purple; fruits dark purple to black. Distribution: *Citharexylum altamiranum* is endemic to Mexico and occurs in pine-oak forests of the Sierra Madres. Though not restricted to one chain of the Sierra Madres, most collections have come from the state of Hidalgo, along the Sierra Madre Oriental. Estimated species: 1 II. *Rehdera* Moldenke, Repert. Spec. Nov. Regni Veg. 39: 48–50. 1935. Type: *Rehdera trinervis* S.F. Blake

## DISCUSSION

### Classification

We present a subgeneric classification of the revised *Citharexylum* including *Baillonia*, which was transferred recently to *Citharexylum* (Christenhusz et al., 2018). Three well-supported, monophyletic subgenera and six sections are recognized. The type species *C. spinosum* is suspected to be of hybrid origin with parental lineages from separate subgenera. This poses a taxonomic problem, since two clades contain representative sequences of this species. However, chloroplast DNA, MSC analyses, fruit morphology, and a predominant distribution in the Caribbean and Florida support the placement of *C. spinosum* in section *Citharexylum* of the Mesoamerican clade (subgenus *Citharexylum*).

### Intergeneric relationships

Phylogenetic reconstructions infer *Rehdera* to be sister to *Citharexylum* and support the inclusion of *Baillonia* in *Citharexylum*. This is consistent with previous studies of Verbenaceae and is bolstered by increased sampling of species in *Citharexylum*. Two species of *Citharexylum, C. oleinum* and *C. tetramerum*, are found to occur among the outgroup (Fig. 3). Preliminary results suggest these species may belong to tribe Duranteae.

As suggested by Marx et al. (2010), *Baillonia amabilis* appears to be a species of *Citharexylum* in which only two of four ovules mature. It is ambiguous as to how the two mature ovules in *Baillonia* are derived (O’Leary et al., 2012). Traditionally, *Baillonia* is described as having one carpel abort early in development, the remaining carpel with two ovules matures. However, dissection of fruits from Rojas 7368 (SI) reveal two carpels, each with one mature ovule. Fruit dissections of closely related species *C. poepiggii* reveal most individuals have four mature ovules, as expected, but one individual, Phillipson 1380 (US), appears to have one abortive carpel (Nataly O’Leary, personal communication). Further investigation of mature carpel/ovule number is needed in *Baillonia* and *Citharexylum* to determine how prevalent abortion of carpels/ovules is throughout the genus.

### Infrageneric relationships

We find support for three major clades of *Citharexylum* that we define as subgenera. The Mexican species *Citharexylum altamiranum* is sister to the rest of the genus, while the remaining species of *Citharexylum* form two major clades defined primarily by geography: (A) a Mesoamerican clade and (B) a South American clade that each contain about half of the taxonomic diversity of *Citharexylum*. We describe *Citharexylum altamiranum*, the Mesoamerican clade, and the South American clade as subgenus *Purpuratum*, subgenus *Citharexylum*, and subgenus *Sudamericanum*, respectively. Each is well defined by a combination of geography and fruit maturation.

The Mesoamerican subgenus *Citharexylum*, comprises three smaller, well-supported clades (clades I, II, and III in Fig. 3), which we describe as sections *Citharexylum, Mexicanum*, and *Pluriflorum*, respectively (Fig. 3). Section *Mexicanum* includes four species of arid-adapted shrubs with small, narrow leaves distributed in the Chihuahuan desert of Central Mexico. Section *Pluriflorum* comprises two species of purple-flowered trees from Central Mexico through Central America. The largest clade within the Mesoamerican clade, section *Citharexylum*, contains 25-30 species of trees and shrubs found in moist to seasonally dry forest. It is a morphologically diverse group with no clear morphological divisions between most of its constituent lineages. Two species are putatively placed in section *Citharexylum*, *C. bourgeauianum* and *C. guatemalense*, which were not included in MSC analyses due to a high proportion of missing data, but nested within this section in gene tree analyses (Appendices S3, S4, S5, S7, and S10). The inclusion of these species is further supported by geography and morphology. *Citharexylum flabellifolium*, an arid-adapted species distributed in Baja California Sur and Sonora, also belongs to this Mesoamerican clade, though its placement within it remains unresolved. Gene trees suggest membership in either section *Pluriflorum* or section *Citharexylum* (Appendices S4 and S7). This species is morphologically distinct from other species in the genus. Like all members of section *Pluriflorum* and some of section *Citharexylum, C. flabellifolium* has purple flowers; however, it differs from the current members of section *Pluriflorum* by its simple, terminal inflorescences that are reduced to 5-15 flowers.

The South American clade, subgenus *Sudamericanum*, consists of three well-supported subclades. Clades IV and V include species with white flowers and red fruits; sect. *Sylvaticum* (clade IV) includes South American trees and shrubs with many-flowered inflorescences that are widely distributed throughout moist forests of South America, including in Amazonia, the Atlantic Coastal Forest of Brazil, and mid elevations of the northern Andes, and sect. *Caribe* (clade V) includes few-flowered shrubs endemic to the Caribbean. Clade VI includes Andean shrubs with a tendency towards producing thorns and black fruits. All members of subgenus *Sudamericanum* have a single fruit color at maturity. *Caribe* and *Sylvaticum* have fruits that are only red at maturity. O’Leary and Frost (in review) identify two morphological groups that correspond to subclades in section *Sylvaticum:* a group of low elevation affiliated species with ternate leaves (e.g., *C. poeppigii* and *C. myrianthum)* and a group of fasciculate-haired species occurring at mid-elevations in the Andes (*C. subflavescens*, *C. montanum*, and *C. kunthianum*). With the exception of *C. amabilis*, a shrub in the flooded lowlands of northern Argentina, southwestern Brazil and Paraguay, all are large trees with white flowers and red fruits. Section *Armatum* (Clade VI), comprises Andean shrub species of *Citharexylum*, including those that are distributed in inter-Andean dry valleys (*C. andinum*, *C. herrerae*, *C. flexuosum*, *C. joergensenii*, *C. peruvianum*, and *C. weberbaueri*), as well as in high elevation cloud forests or grasslands (*C. argutedentatum*, *C. dentatum*, *C. ilicifolium*, *C. pachyphyllum*, *C. reticulatum*, and *C. sulcatum*). One additional species in subgenus *Southamericanum* is tentatively placed in section *Sylvaticum*. *Citharexylum reticulatum* had too much missing data to include the MSC analysis, but was strongly supported as being related to the other members of section *Sylvacticum*. This species would be difficult to place by morphology alone: it has the long, many-flowered inflorescences with white flowers that are typical of section *Sylvaticum*, but differs in many ways, including its small stature, reportedly black fruits (Moldenke, 1957-1959), and its distribution at higher elevations, all of which would be consistent with its placement in section *Armatum*, the Andean shrub clade.

Section *Armatum* is a particularly difficult group to parse further taxonomically. Its species all have synchronous fruit maturation, but the ripe fruit color can be either red or black, and fruit color does not define lineages within this section. Similarly, many species of this section also produce axillary shoots that are modified into thorns— the first character applied in the history of classification of *Citharexylum*; however, thorny species do not form a clade. While monophyly of the section and major lineages within section *Armatum* are strongly supported, the relationships between those lineages remain unresolved. Weak support (bootstrap values ≤ 70, posterior probability ≤ 95%) and short branch lengths between them suggest rapid divergence between the three lineages (Figs. 3 (support only; Appendix S12 for branch lengths) and 4). Supporting the hypothesis that sect. *Armatum* represents a rapid radiation, species share morphological traits in a variety of combinations that suggest potential sorting of ancestral variation within a short time span (Pease et al., 2016). The section’s geographic distribution also predicts rapid diversification, which is well documented among Andean plant lineages (Hughes and Eastwood, 2006; Peret et al., 2012; Madriñán et al., 2013; Luebert and Weigend, 2014; Uribe-Convers and Tank, 2015; Lagomarsino et al., 2016; Diazgranados and Barber, 2017; Vargas et al., 2017).

Sampling of Caribbean species is relatively low in this study; five out of ten described species were sampled (*C. alainii, C. caudatum, C. ellipticum, C. microphyllum*, and *C. spinosum*). Section *Caribe* contains two species, *C. alainii* and *C. microphyllum*, which are Caribbean endemics. The other three species, *C. caudatum, C. ellpiticum*, and *C. spinosum*, are placed in in section *Citharexylum* and are widespread coastal species with ranges that extend into the Caribbean from Mesoamerica and/or South America. The unsampled species, *C. albicaule, C. discolor, C. schulzii, C. stenophyllum*, and *C. tristachyum*, are Caribbean endemics. Inclusion of these species is needed in future studies to determine if they belong in section *Caribe* or elsewhere in the updated classification scheme.

### Fruit evolution

*Citharexylum* (including *Baillonia*) represents an independent origin of fleshy fruits in Verbenaceae; its ancestors likely had schizocarps, like those produced by *Rehdera*, sister to the rest of Citharexyleae (O’Leary et al., 2012). Within *Citharexylum*, there has been dynamic fruit color evolution. Mature fruits that are black (e.g., Fig 1L) have evolved multiple times independently from red-fruited ancestors (e.g., Fig. 1K), including in the species sister to the rest of the genus (*C. altamiranum*), at least once within the Andean shrubs of section *Armatum*, and on various occasions in subgenus *Citharexylum*. Within the Mesoamerican subgenus *Citharexylum*, black fruits are always associated with sequential maturation in which fruits mature from green to red-orange to finally black, with both colors of fruits observed simultaneously in infructescences (Fig 1J). According to ancestral state reconstructions, sequential maturation has either evolved twice within the subgenus—once in the common ancestor of section *Pluriflorum* and again in the common ancestor of section *Citharexylum*—or evolved once in the common ancestor of *C*. subg. *Citharexylum* and lost multiple times (Appendix S15). Reversals from sequential maturation to a single mature fruit color need to be confirmed via field observation, since it is unclear if fruits of those species mature at the red-orange stage, or if observations for fully mature black fruits are lacking. It is likely that fruit color plays an important role in the attraction of bird seed dispersers of *Citharexylum*. Both red and black fruits are known signals to bird dispersers (Wheelwright and Janson, 1985). Further, sequential maturation process is well-documented in other plant groups (Sinnott-Armstrong et al., 2018; Sinnott-Armstrong, 2020). A recent hypothesis posits that red fruits from black fruited ancestors are a result of paedomorphosis (Box and Glover, 2010), and perhaps represent an adaptation to shorter growing seasons in northern temperate regions (Sinnot-Armstrong et al 2018). This specific hypothesis is not supported in the exclusively tropical *Citharexylum*, nor does there seem to be a connection between seasonality in dry habitats and the loss of sequential maturation in subgenus *Citharexylum*. Fruit color could also be connected to the degree to which a plant species is able to invest in costly anthocyanins; darker colors of fleshy fruits are perceived as an honest signal of food quality in birds, and thus would be beneficial in conditions that allow this costly investment (Schaefer et al., 2008).

**Fig. 1.**
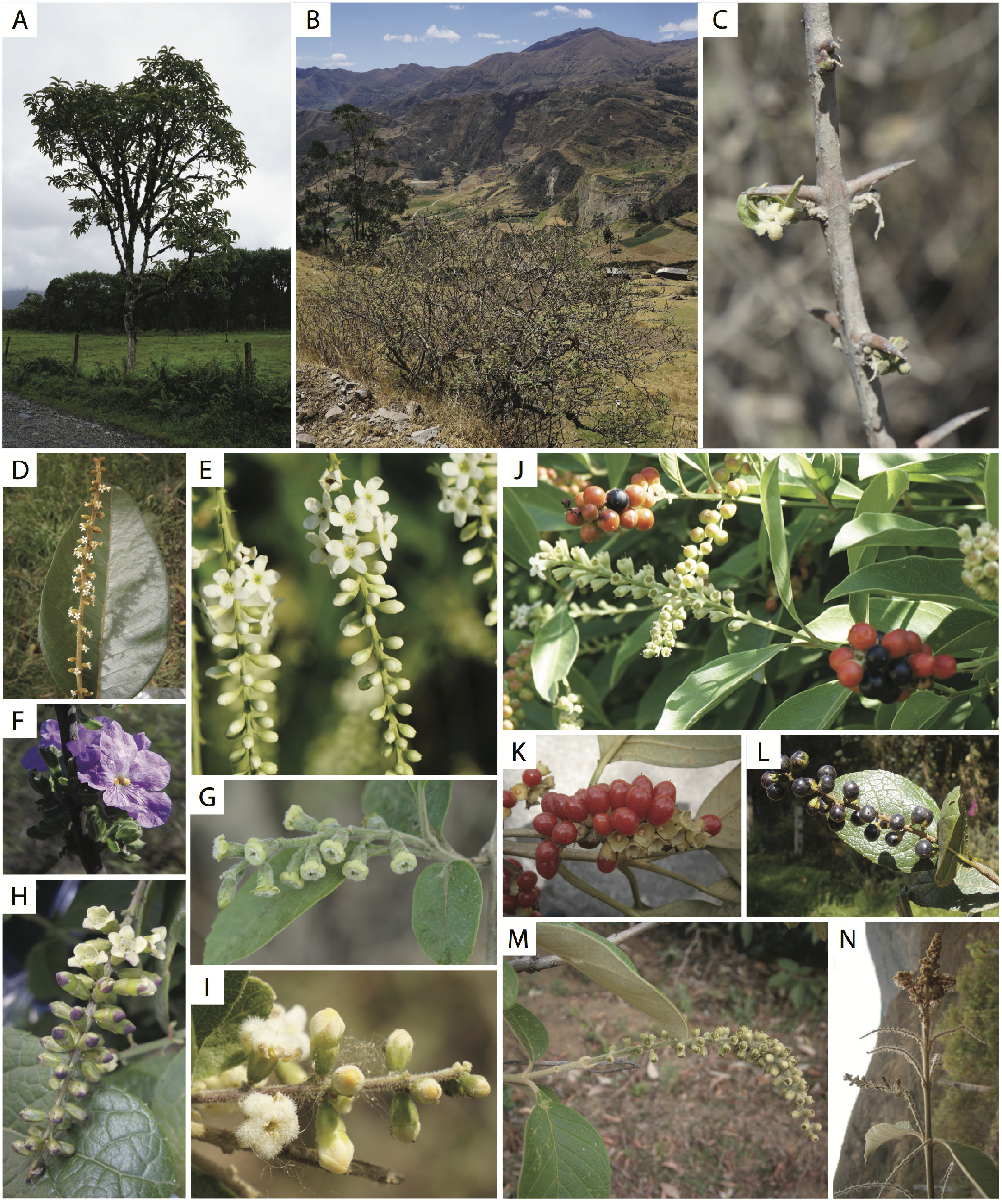
Morphological diversity of *Citharexylum*. Habit (A-B): (A) tree (*C. montanum*), (B) shrub (*C. weberbaueri*); Thorns (C; *C. flexuosum*); Flower color (D-I): (D, E) white (*C. subflavescens, C. hexangulare*), (F) purple (*C. flabellifolium*), (G) pale yellow-green (*C. kobuskianum*), (H) pale with purple-tipped buds (*C. sulcatum*), (I) white with orange-tipped buds (*C. peruvianum*); Fruit color (J-L): (J) sequential maturation from green to orange to black (*C. ellipticum*), (K) synchronous maturation, green to red (*C. montanum*), (L) synchrounous maturation, green to black (*C. sulcatum*); Inflorescence (M-N; *C. kunthianum*): (M) pistillate inflorescence with simple terminal raceme (N) staminate infloresence with compound terminal racemes and simple axillary racemes. All photos by L. Frost, except for the photo of *C. flabellifolium* in F, which was taken by Carianne Campbell and used here with the permission of the creator.

**Fig. 2.**
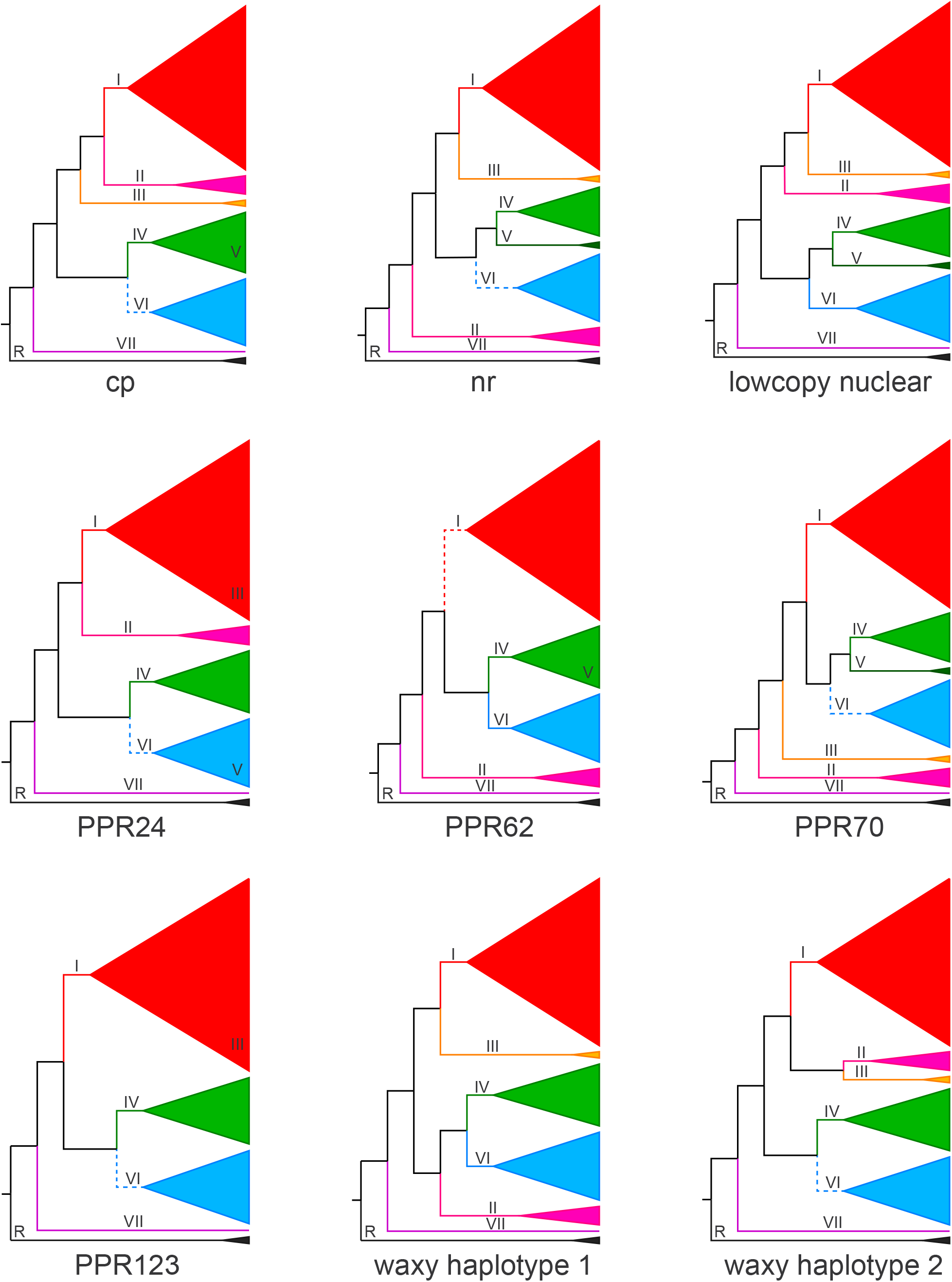
Gene tree cartoons for cp, nr, lowcopy nuclear combined, and individual lowcopy nuclear datasets showing topological differences among gene trees. Roman numerals on branches correspond to clades of the tree recognized in the text; R marks the genus *Rehdera*. For gene trees in which one clade nested within another, the numeral for that clade is placed within the inclusive clade (e.g. clade III in PPR24 and PPR123). Data for species in clade II were missing for PPR123, for clade III for PPR62, and for clade V for PPR123 and both waxy haplotypes. Solid lines on branches indicate clades; dashed lines indicate grades. Tip labels may be found in Fig. 3.

**Fig. 3.**
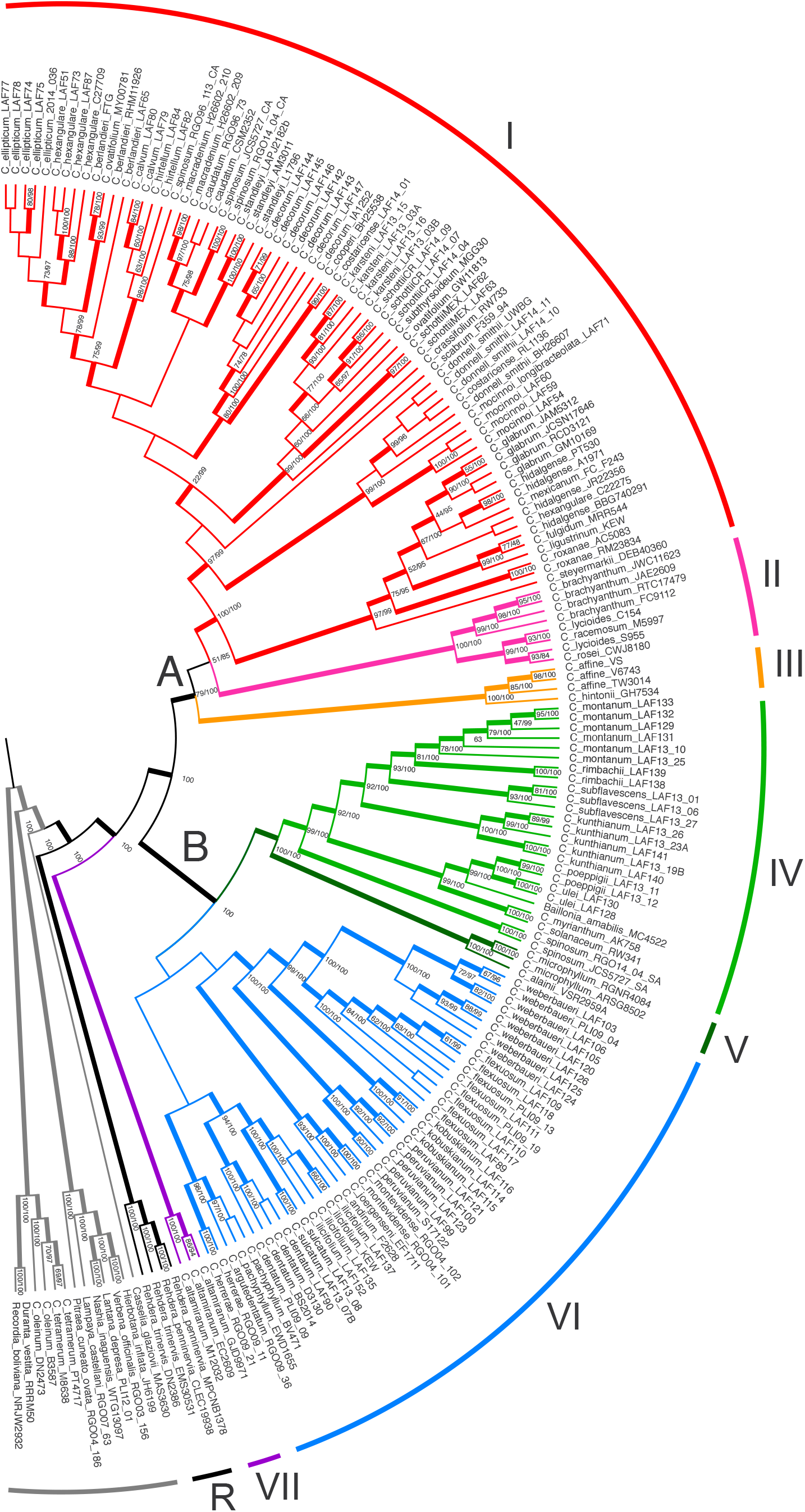
Cladogram of Bayesian consensus tree for all data combined. Thickened branches indicate high support (≥ 70 bootstrap support from maximum likelihood (ML) analyses or ≥ 90 % posterior probability from Bayesian inference (BI)). Support for nodes are plotted as bootstrap/posterior probability (percent). Roman numerals, curved lines at the tips, and branch colors correspond to clades recognized in the text; Clades A and B are also referred to as the Mesoamerican and the South American clade, respectively. R indicates the genus *Rehdera*.

**Fig. 4.**
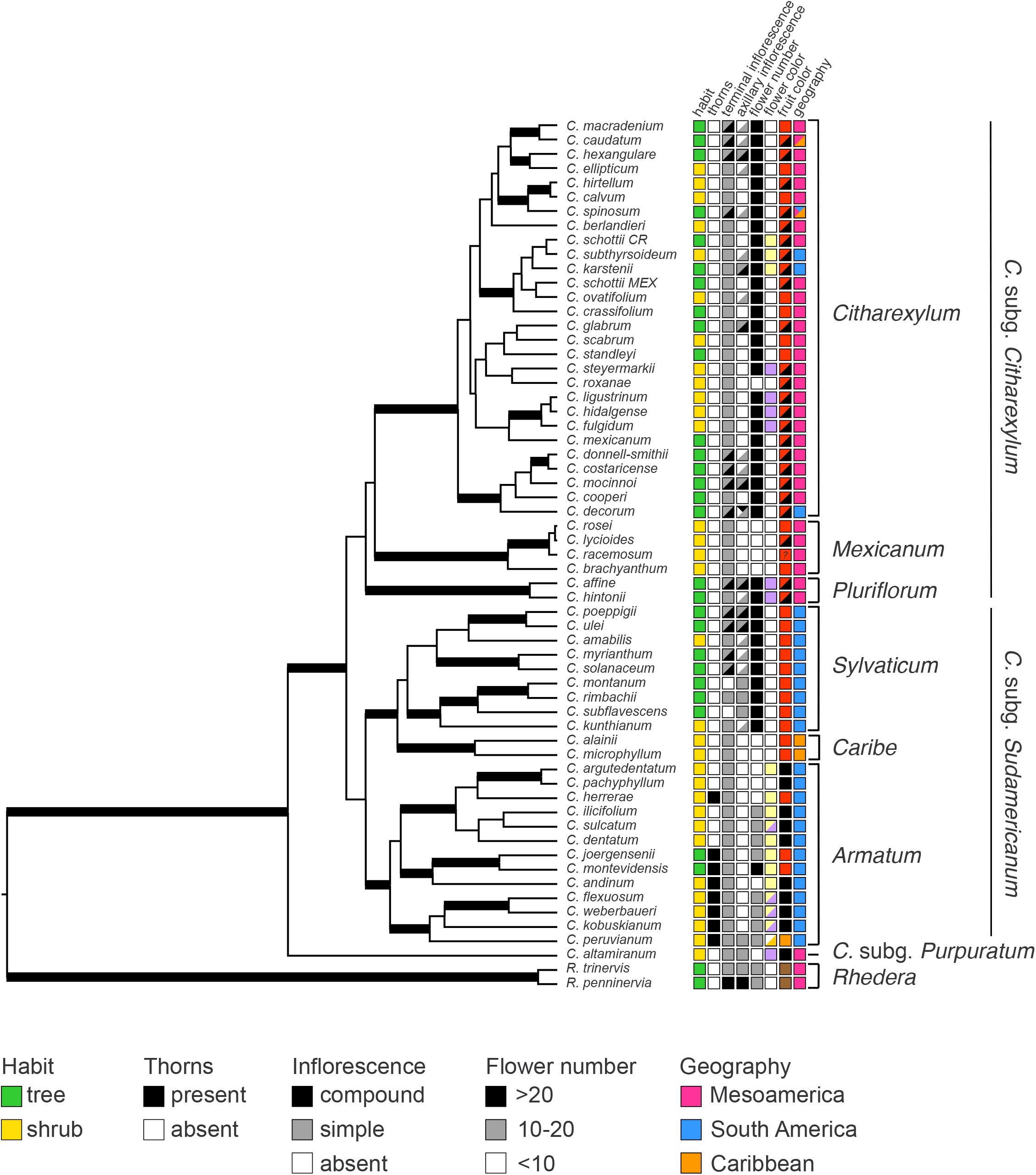
BEAST multi-species coalescent tree. Thickened branches indicated nodes with high support (posterior probability ≥ 0.95). Colored boxes represent character states of species. First column from the left—habit: tree (green) or shrub (yellow); thorns: present (black) or absent (white); terminal inflorescence: compound (black), simple (gray), or absent (white); axillary inflorescence: compound (black), simple (gray), or absent (white); flower number: >20 (black), 10-20 (gray), or <10 (white); flower color: literal (white, pale yellow, purple, pale yellow with purple lobes (split yellow and purple boxes), white with orange lobes (split white and orange box)); mature fruit color: literal (red with no intermediate, black with no intermediate, black with red/orange intermediate (split orange and black boxes)), or dry fruits (brown); Geography: Mesoamerica (pink), Caribbean (orange), or South America (blue), boxes with multiple color inhabit multiple regions. The classification scheme of *Citharexylum* corresponds to clades identified in previous figures: *C*. subg. *Citharexylum* (A; also referred to as the Mesoamerican clade), *C*. subg. *Sudamericanum* (B; also referred to as the South American clade), section *Citharexylum* (I), section *Mexicanum* (II), section *Pluriflorum* (III), section *Sylvaticum* (IV), section *Caribe* (V), section *Armatum* (VI), and section *Purpuratum* (VII). R indicates the genus *Rehdera*.

### Flower color evolution

Purple floral pigmentation has arisen at least four times within *Citharexylum* (Fig. S6). This includes in the common ancestor of section *Pluriflorum* and subgenus *Purpuratum*, twice within each section *Citharexlyum* and *Armatum*, in which the flower buds of several species are tipped with purple. These species also have dark purple to black fruits. Though not all black-fruited species have pigmented flowers, there may be some connection between fruit color and expression of anthocyanins in the flowers, as has been found in some plant communities (Renoult et al., 2014). Species with black fruits also occur at high elevations in the Andes, where UV radiation is higher (Steyn, 2008). In these species, darker pigments in flowers and fruits might offer some protection to developing reproductive structures and seeds by absorbing excess UV radiation. This hypothesis does not apply, however, to the purple flowered species in Mesoamerica, which do not occur at high elevations. Flower color in these species may be linked to other factors such as pollination, but little is known about this aspect of their natural history. However, the large, showy corollas of some purple flowers (e.g., Fig. 1F) provide an ideal starting point to further understand the ecological role of flower color in pollination of *Citharexylum*.

### Infloresence evolution and dioecy

Dioecy needs further investigation throughout *Cithareyxlum*. Several species have been confirmed as dioecious in the literature, including *C. spinosum* (Tomlinson and Fawcett, 1972), *C. myrianthum* (Rocca and Sazima, 2006), and all Costa Rican species (Rueda and Hammil, in prep.). Dioecy has been observed in the field for many others (L. Frost, personal observation); it can be inferred in *Citharexylum* from observation of post-anthesis inflorescences. In staminate individuals, the entire flower disarticulates from the pedicel at senescence, leaving naked racemes. In pistillate individuals, the calyx becomes lignified as the fruits mature and is retained on the pedicel, even as fruits are eaten or fall off. This pattern is observed in specimens of *Baillonia* (see Palmer 1853-6 [Orrell and Howell, 2018] and Hassler 2683a [MNHN, 2018a] for putative staminate and Prance et al. 15 [Tulig et al., 2017] and Weddell 3193 [MHNH, 2018b] for putative pistillate specimens), but more detailed study is needed to determine if this species is dioecious.

Variation in inflorescence architecture may additionally support our hypothesis that dioecy is more widespread than has been confirmed in the literature. Functionally staminate plants often have greater investment in production of attractive, reproductive structures, which can translate to a greater number of flowers produced overall than pistillate individuals (Mayer and Charlesworth, 1991). Variation within species of *Citharexylum* for presence/absence of compound or axillary inflorescences may be related to sex. In these variable species, staminate individuals could increase flower production either through the development of axillary inflorescences, or compound inflorescences, or both. Variation between sexes is easily observed for species with large, many-flowered inflorescences, but is difficult to detect in shrub species with reduced inflorescences. These species have simple, few-flowered inflorescences that terminate short, axillary branches. One individual does not obviously produce more flowers or more inflorescences than another. It is also more difficult in general to infer the sex of individuals in these few-flowered species using the observations described above and dioecy has not been postulated for any of these species. Further investigation of dioecy in the Andean and arid-adapted Mexican shrubs is warranted.

Cryptic dieocy is not always readily observable and may be more widespread in the family. A hand-written note on the label of G. Diggs & M. Nee 2473 (F) *C. oleinum* specimen reads, “4 epipetalous, ± sessile stamens. Style short, functionally a male flower?” *Citharexylum* is the only genus in Verbenaceae in which dioecy has been documented. Citharexylum *oleinum* and *C. tetramerum*, which consistently fell outside of Ctiharexylum in phylogenetic analyses, would represent a separate origin in Verbenaceae. The only other identified occurrence of dioecy in Verbenaceae is in *Lippia* section *Dioicolippia* (Múlgura, 2010), although gynodioecy also has been documented in *Rhaphithamnus venustus* (Crawford et al., 1993).

### Evolution of dry-adaptive traits

Of the morphological traits examined, habit, axillary inflorescences, and flower number were found to be significantly correlated with each other. In *Citharexylum* combinations of these traits were typically found in shrubs with simple, exclusively terminal inflorescences (i.e., no axillary inflorescences) and few flowers per inflorescence, or trees with many-flowered inflorescences that often include axillary inflorescences, compound inflorescences or both. These alternate suites of traits are found repeatedly across the phylogeny of *Citharexylum*. For example, shrubs with few-flowered, simple, terminal racemes occur in at least four distantly related lineages: *C. roxanae*, *C. flabellifolium* (phylogenetic placement uncertain), section *Armatum*, section *Caribe*, and section *Mexicanum*. While not closely related, these clades share similar environmental constraints: limited water availability in their high elevation and/or arid to semi-arid biomes. These environmental conditions are known to constrain above-ground growth (Shantz, 1927), and woody plants are often reduced to shrubs in these habitats. Plants growing at high elevations may be additionally subject to high winds, cloud cover, and low temperatures, which can also limit the height and reproductive effort of woody species (Howard, 1970). We suggest that this combination of characters— short habit and reduced investment in reproductive growth— also reflects a high/dry habitat syndrome in *Citharexylum*.

This syndrome is sometimes associated with the evolution of armament. While there are several possible reconstructions of the evolutionary history of thorns (Figure SX) due to lack of phylogenetic resolution in section *Armatum* (see discussion above), it seems likely that there are at least two origins of thorns (Fig. S2; SX). These may have evolved as a defense mechanism against large herbivores in the more open vegetation of inter-Andean dry valleys. This reflects convergent evolution with other members of the flora in this stressful environment that are also characterized by thorn production (Mazancourt and Loreau, 2000). Thorns were retained in the more mesic, lowland tree species *C. montevidense*, which dispersed to the gallery forests in the savannas of southern Brazil, eastern Argentina, Paraguay, and Uruguay from within an otherwise Inter-Andean Valley clade. An independent origin of thorns is strongly supported for *C. herrerae*, which is more closely related to the high-elevation grassland/cloud forest species that lack thorns. Though not technically bearing thorns, some species outside of the inter-Andean dry valley shrubs also have sharp defenses. *Citharexylum amabilis* has small, sharp projections borne out of leaf scars (referred to a “spiny sterigmata” by Moldenke, 1941), and many arid-adapted Mexican shrubs (*C. brachyanthum*, *C. flabeillifolium*, *C. lycioides*, *C. racemosum*, and *C. rosei*) have relatively short, pointed, branches (Moldenke 1957-1959).

The evolution of distinctive leaf traits is also associated with the dry-adaptive syndrome in *Citharexylum*. Though they share some traits related to adaptation to arid environments *C. roxanae*, *C. flabellifolium*, and section *Mexicanum* have evolved different vegetative strategies for coping with life in the desert. Whereas many Andean species of section *Armatus* have tough, leathery leaves along with their thorns, *Citharexylum roxanae* has chartaceous, ephemeral leaves and an ephedroid habit in which the persistently green stems likely contribute to photosynthesis when most of the leaves have fallen. *Cithareyxlum flabellifolium*, which occurs with *C. roxanae* in Baja California Sur, but extends further into Sonora, has fan-shaped leaves that are densely hairy, presumably protecting leaf tissue from the sun and maintaining humidity around stomatal pores (Ehleringer, 1982). Finally, the four species of section *Mexicanum* found in the Chihuahuan desert have small, narrow leaves that dissipate heat better than the large, broad leaves that characterize other subclades, reducing the need for evaporative cooling and water loss from the process (Solbrig and Orians, 1977).

The remaining dry-adapted shrub clade, section *Caribe*, consists of species inhabiting limestone outcroppings of semi-arid slopes in the Caribbean. These species exhibit a mixture of the aforementioned leaf traits associated with more arid environments: *C. alainii* has somewhat larger, subcoriaceous leaves (as does *C. albicaule*, an unsampled Caribbean endemic), whereas *C. microphyllum* has small, slender leaves (as does *C. stenophyllum*). Finally, *C. schulzii*, another unsampled Caribbean endemic that may belong to section *Caribe*, has small, coriaceous leaves.

### Convergent evolution

The multiple origins of each of the studied traits underscores the broader difficulty in describing diversity in Citharexyleae. In general, clades lack synapomorphies that evolved a single time, and instead sections can be defined based on combinations of multiple traits, including habit, inflorescence architecture, fruit color, and geographic distribution. Other groups, like section *Citharexylum*, lack even a series of unifying traits, and instead contain as much morphological diversity as is present across the entire genus. This labile morphological trait evolution has hampered efforts at classification within *Citharexyleae*, however, it is incredibly interesting from an evolutionary perspective. For example, convergence towards a suite of traits associated with multiple colonizations of stressful, often arid environments from ancestors in mesic habitats has been demonstrated. Further, traits that are associated with ecological relationships, such as fruit and flower color, also have complex histories that may reflect local adaptation to different pollinator and disperser pools. As we demonstrated with dry-adapted species, diversity across the genus is often linked to climatic factors and has evolved convergently in similar habitats in Mesoamerica and South America. Future work is aimed at understanding patterns of movement and evolution into (and out of) biomes that led to the present distribution of *Citharexylum* and likely drove the repeated evolution of suites of environmentally-relevant characters.

### Conclusions

We present the first phylogeny of Citharexyleae to include near complete taxonomic sampling, confirming earlier hypotheses of intergeneric relationships. The first robust phylogeny of *Citharexylum* provides a framework to identify major clades, map morphological characters which define those clades, and understand patterns of character evolution. Three subgenera are described based on fruit maturation and geography. Six sections are described based on habit and inflorescence architecture, some by flower color and fruit color as well. Morphology at lower ranks, sections and subclades often corresponds with environment. *Citharexylum* is a widespread in the Neotropics and inhabits multiple biomes. We demonstrate a dynamic pattern of trait evolution, including the correlated evolution of multiple traits associated with colonization of dry habitats. This labile trait evolution likely facilitated the rich biogeographic history of the genus, reflected here in the morphological diversity. Future studies will examine in detail the biogeographic patterns underlying diversity in *Citharexylum*, as well as continue to clarify evolutionary relationships and refine taxonomic limits in parts of *Citharexylum* characterized by rapid radiations.

## ACKNOWLEDGEMENTS

The authors thank the curators of the following herbaria for permitting us to sample specimens: F, A/GH, MEX, MO, SI, TEX, USM, WTU; the University of California Botanical Garden at Berkeley for contributing silica-dried tissue of *C. hidalgense;* the following people for assistance in the field: Carlos Burelo-Ramos, Warren Cardinal-McTeague, Itzue Caviedes-Solis, Ross Furbush, David Garcia, Barry Hammel, Johan Home, Diego Morales-Briones, Lenis Prado, Sarah Tyson, and Simon Uribe-Convers; and anonymous reviewers. This study was supported by NSF grants DEB 0542493, DEB 0710026, and DEB 1020369 to RGO and DEB 1500919 to LAF and RGO.

## AUTHOR CONTRIBUTIONS

L.F., D.T., and R.O. designed the research. L.F. and R.O. conducted fieldwork. L.F. collected and analyzed data, and led the writing with significant contributions from the other authors. N.O. provided species identifications, morphological data, and revised the classification scheme. R.O., N.O., and L.L. critically revised the manuscript.

## DATA AVAILABILITY STATEMENT

Submission of Illumina MiSeq reads to the NCBI Sequence Read Archive (SRA) is in progress under submission number SUB7989566 and will be made available upon publication. All other data formats (tree files, alignments, character matrices, etc.) will be uploaded on dryad and made available upon publication.

## SUPPORTING INFORMATION

Additional Supporting Information may be found online in the supporting information section at the end of the article.

**Appendix S.1.**
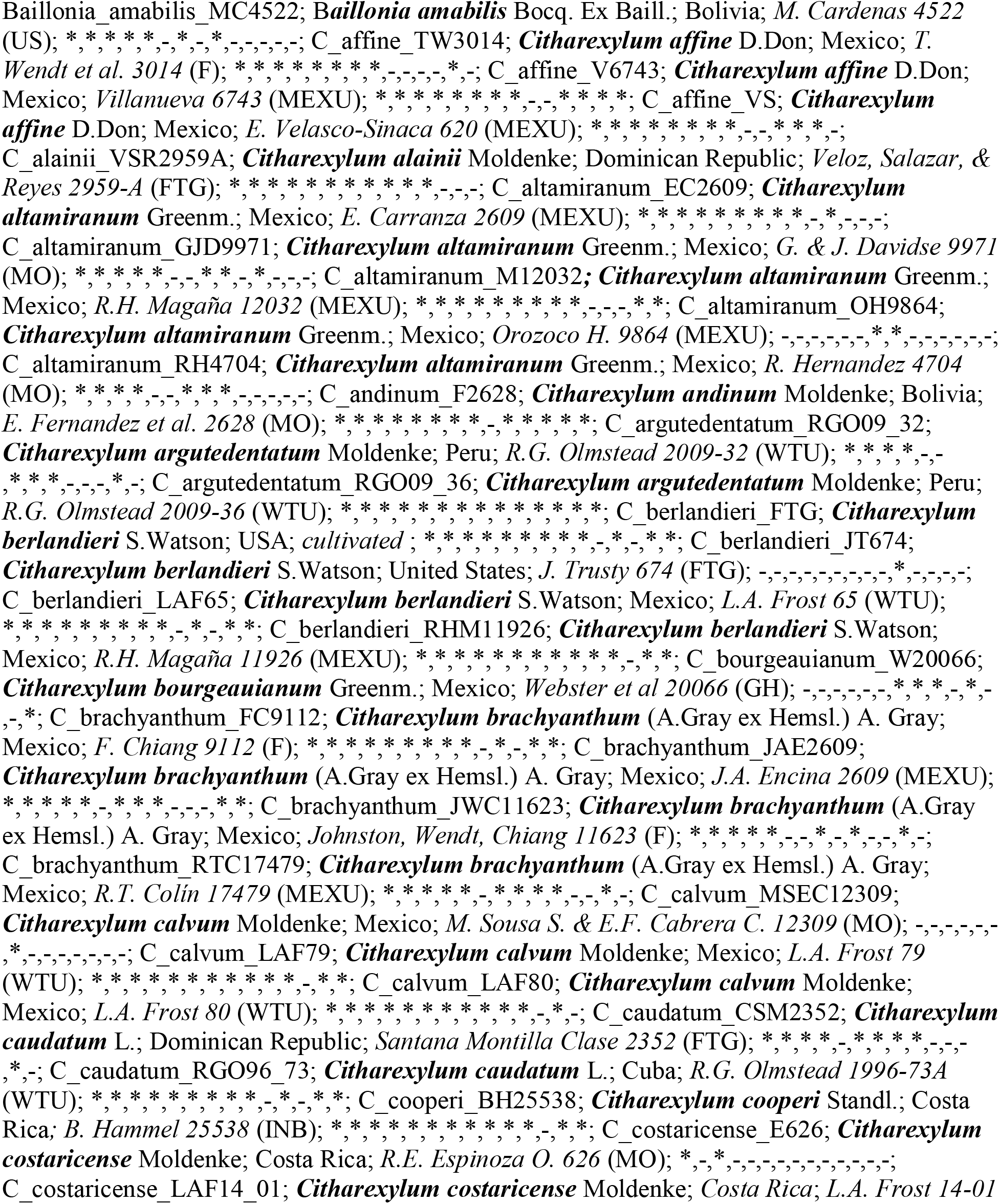
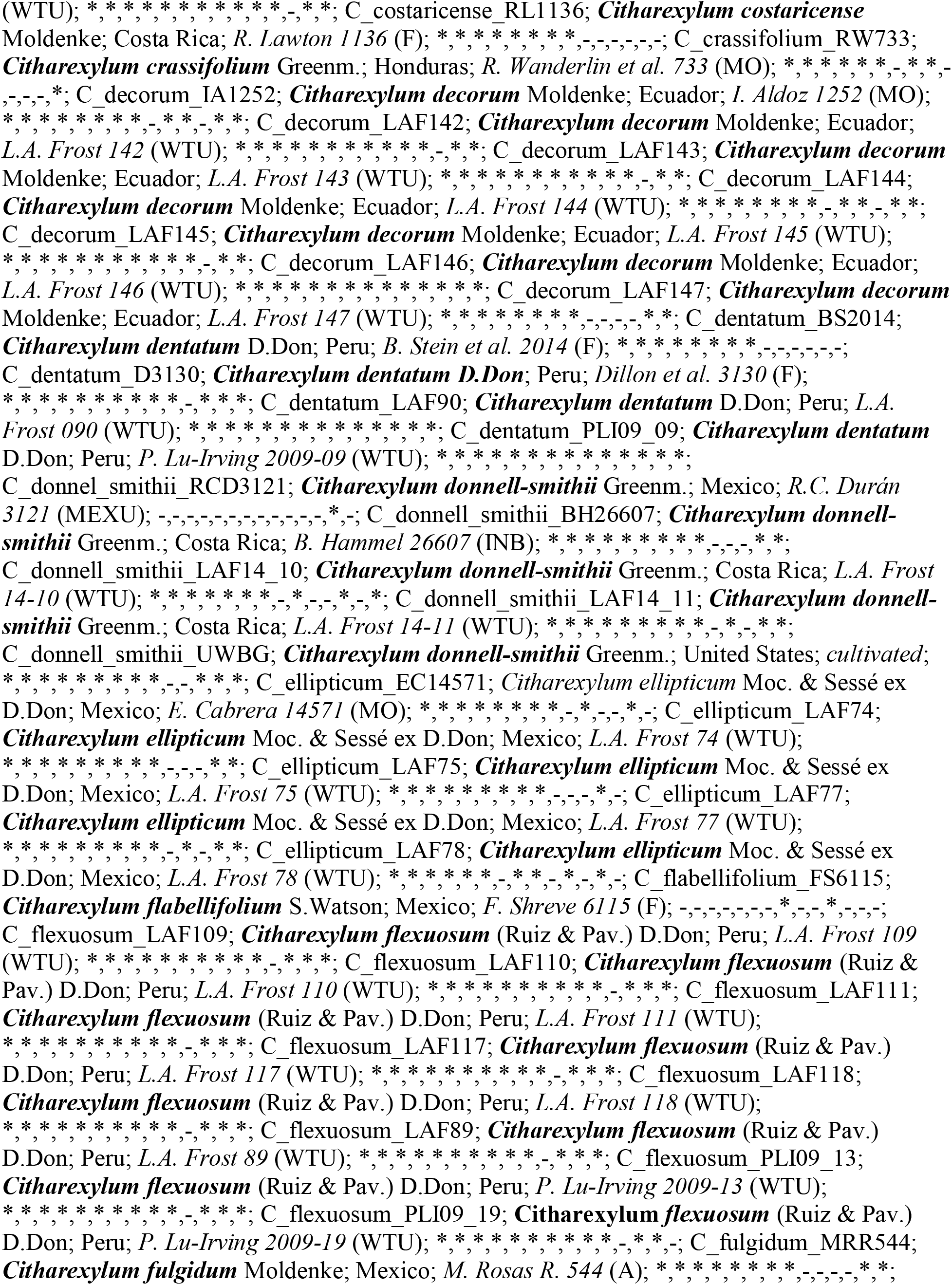
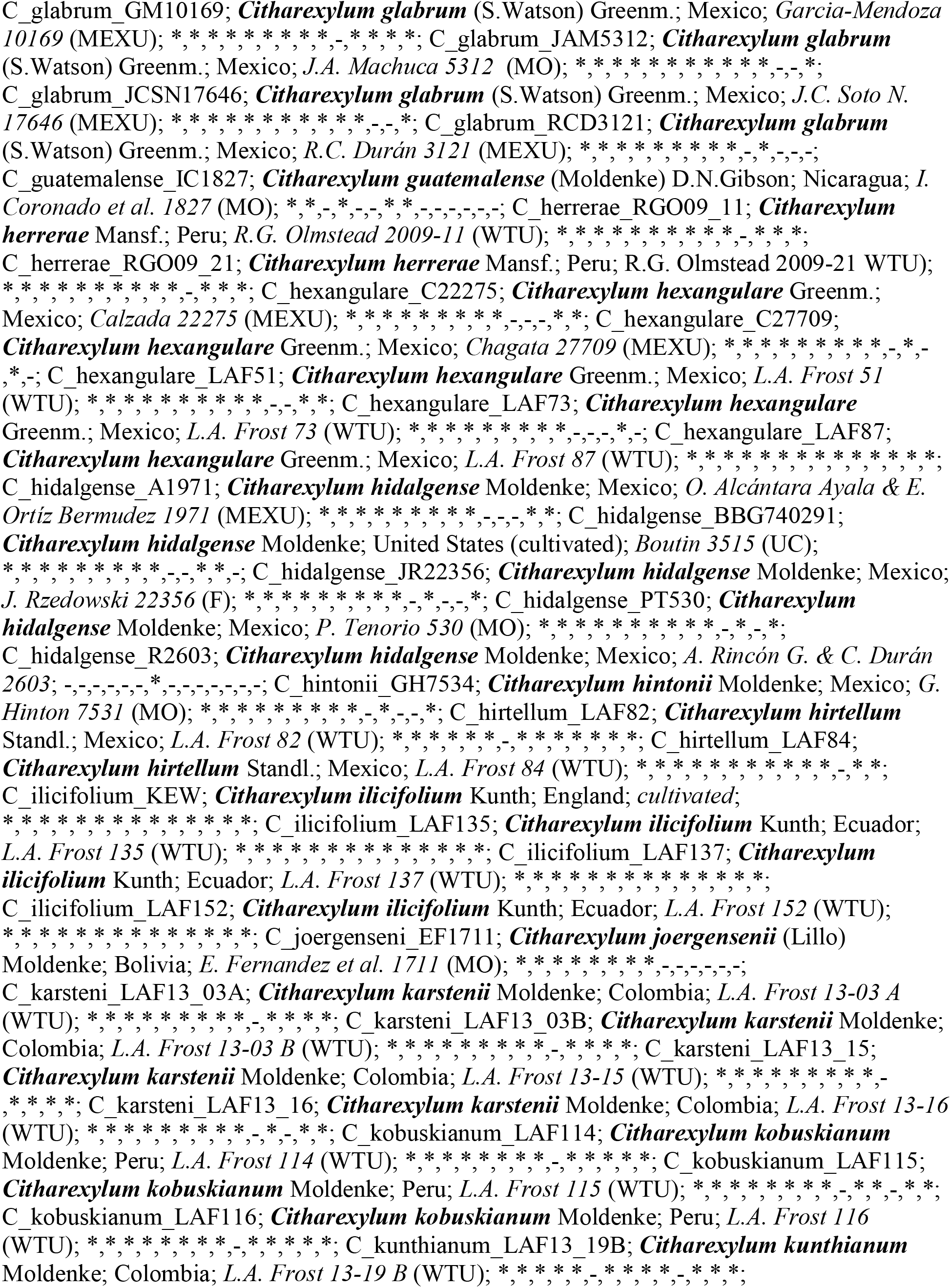
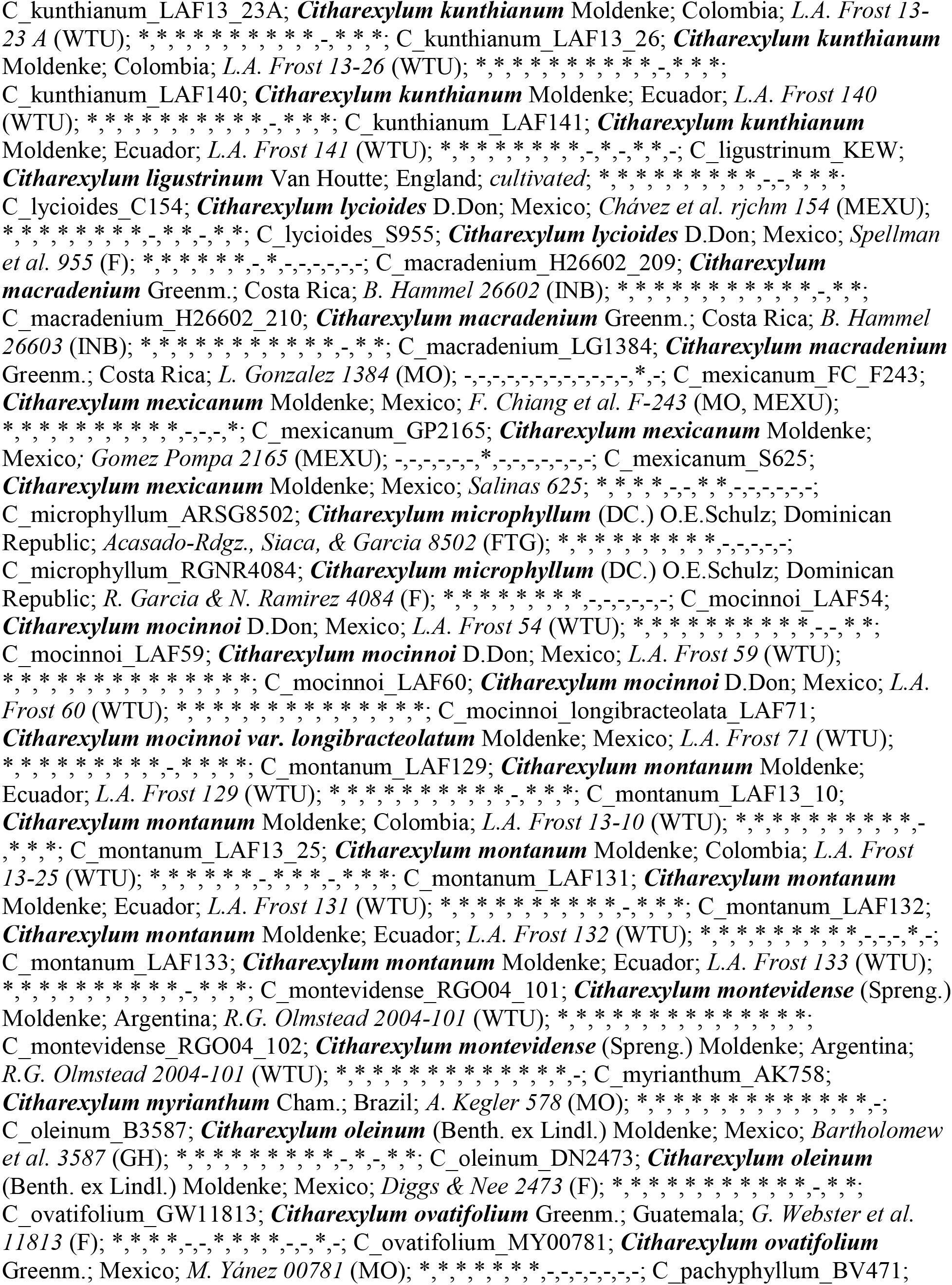
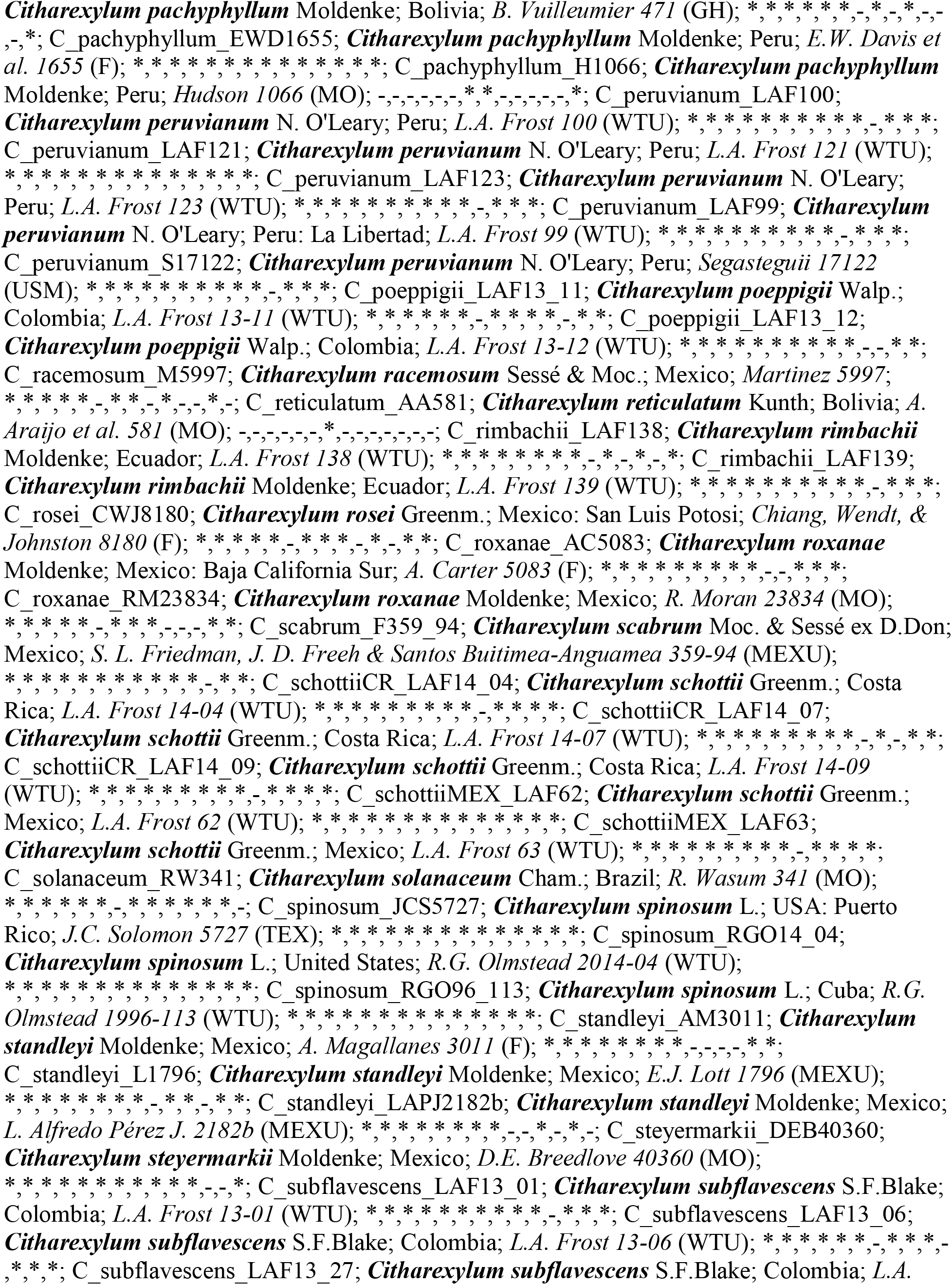
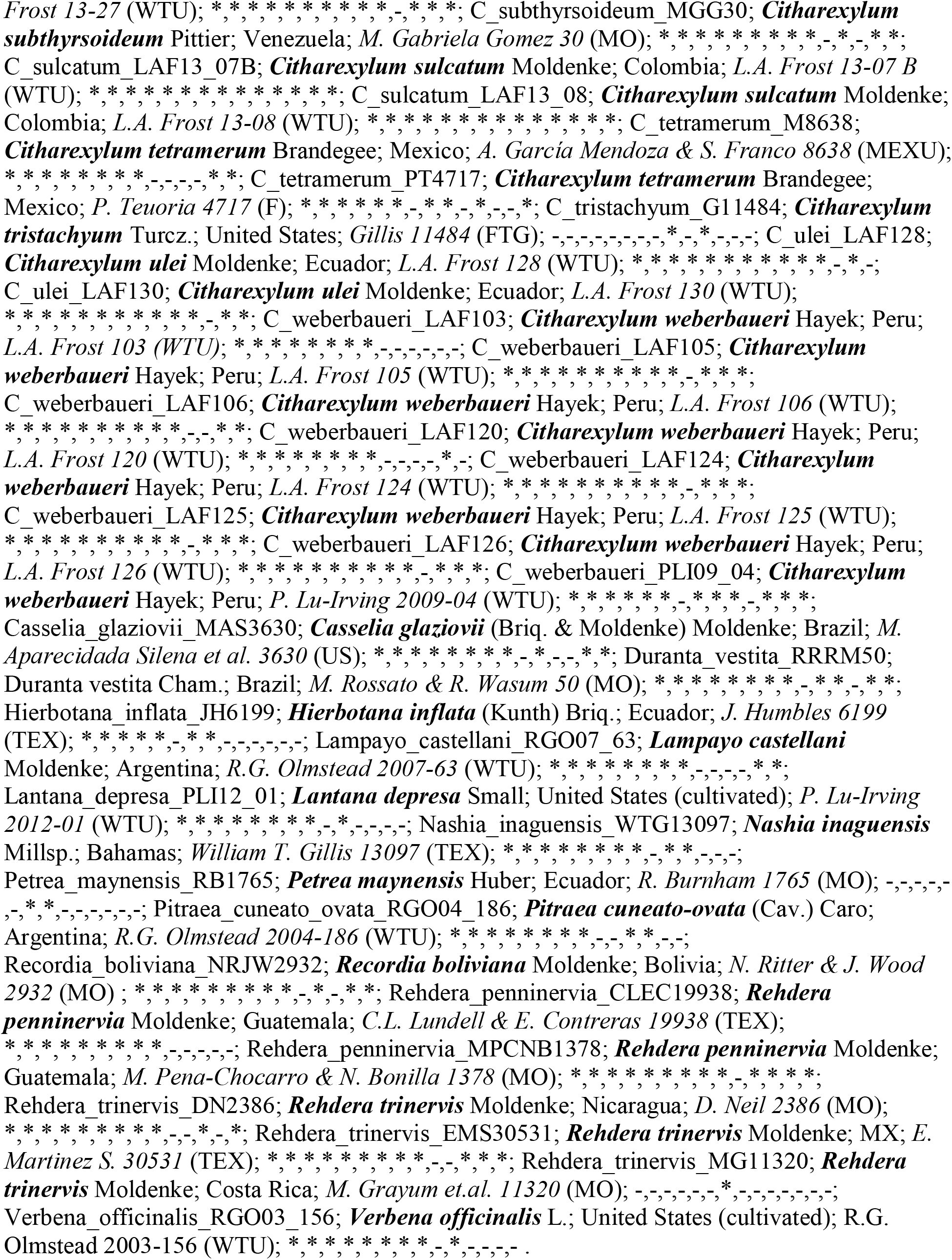
Sequence ID; Taxon (accepted); Locality; Voucher (Herbarium); Sequence data: ndhF, rpoC2, trnT-L-F, rbcL, matK, rpl32, ETS, ITS, PPR24, PPR62, PPR70, PPR123, waxy haplotype 1, waxy haplotype 2; hyphens indicate missing sequences and asterisks indicate newly obtained sequences for this study.

**Appendix S.2.**
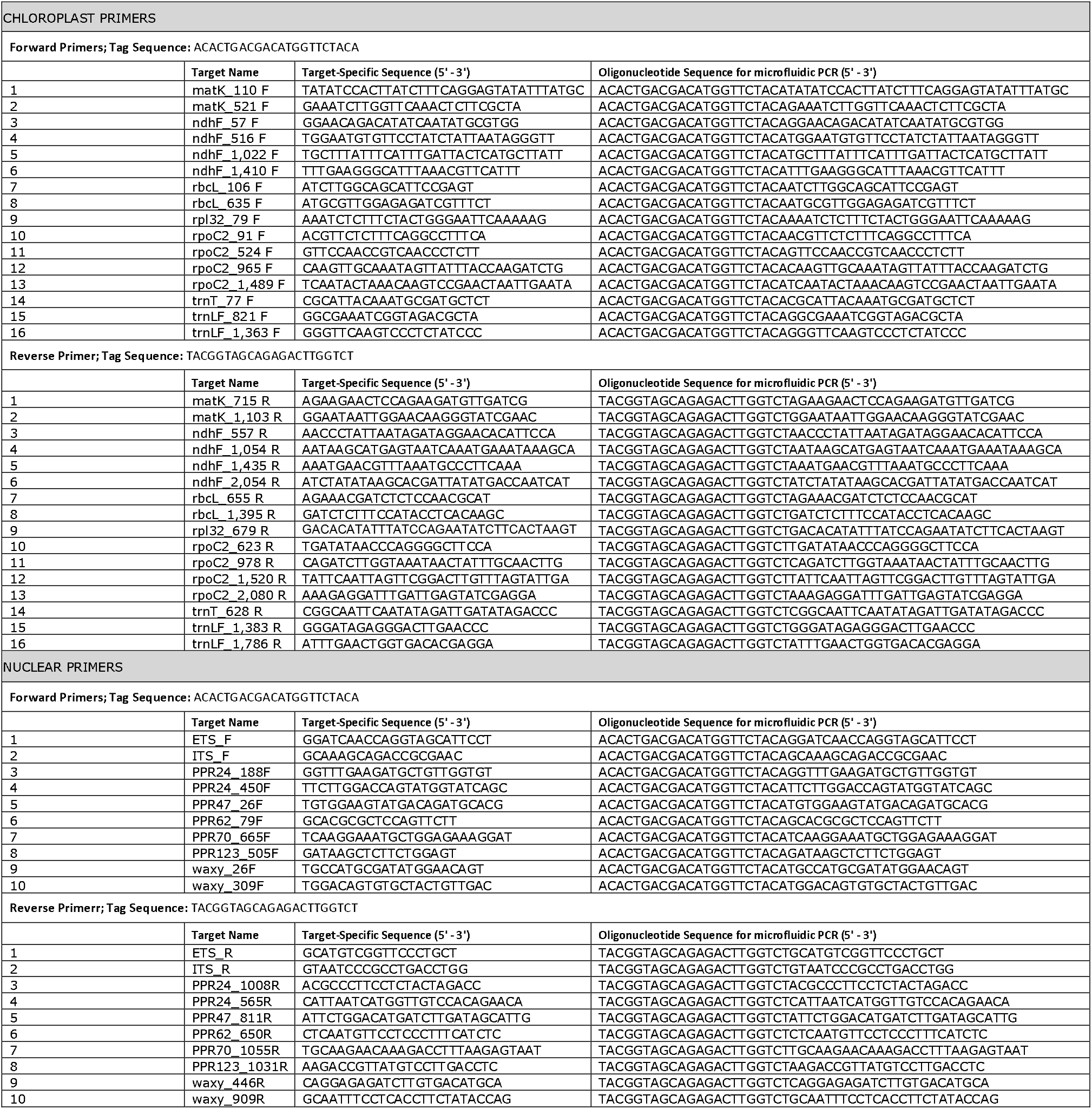
Primer pair sequences.

**Appendix S.3.** Gene trees for cpDNA. Right: maximum likelihood (ML) tree from RAxML; left: Bayesian consensus tree (BI) from MrBayes.

**Appendix S.4.** Gene trees for nrDNA. Right: maximum likelihood (ML) tree from RAxML; left: Bayesian consensus tree (BI) from MrBayes.

**Appendix S.5.** Gene trees for PPR24. Right: maximum likelihood (ML) tree from RAxML; left: Bayesian consensus tree (BI) from MrBayes.

**Appendix S.6.** Gene trees for PPR62. Right: maximum likelihood (ML) tree from RAxML; left: Bayesian consensus tree (BI) from MrBayes.

**Appendix S.7.** Gene trees for PPR70. Right: maximum likelihood (ML) tree from RAxML; left: Bayesian consensus tree (BI) from MrBayes.

**Appendix S.8.** Gene trees for PPR123. Right: maximum likelihood (ML) tree from RAxML; left: Bayesian consensus tree (BI) from MrBayes.

**Appendix S.9.** Gene trees for waxy haplotype 1. Right: maximum likelihood (ML) tree from RAxML; left: Bayesian consensus tree (BI) from MrBayes.

**Appendix S.10.** Gene trees for waxy haplotype 2. Right: maximum likelihood (ML) tree from RAxML; left: Bayesian consensus tree (BI) from MrBayes.

**Appendix S.11.** Gene trees for low-copy nuclear genes combined. Right: maximum likelihood (ML) tree from RAxML; left: Bayesian consensus tree (BI) from MrBayes.

**Appendix S.12.** Gene trees for all loci combined. Right: maximum likelihood (ML) tree from RAxML; left: Bayesian consensus tree (BI) from MrBayes.

**Appendix S.13.** (A) Maximum likelihood and (B) maximum parsimony ancestral character reconstructions for habit. Pie charts at nodes in A indicate the probability of a given state at that node—tree (green) or shrub (yellow). Letters in boxes at nodes in B indicate the state(s) reconstructed at that node—T=tree and S=shrub—coloration on branches indicates the state(s) reconstructed along the branch and matches the color scheme in A.

**Appendix S.14.** (A) Maximum likelihood and (B) maximum parsimony ancestral character reconstructions for the presence/absence of terminal inflorescences. Pie charts at nodes in A indicate the probability of a given state at that node—present (black) or absent (white). Letters in boxes at nodes in B indicate the state(s) reconstructed at that node—A=absent and P=present—coloration on branches indicates the state(s) reconstructed along the branch and matches the color scheme in A.

**Appendix S.15.** (A) Maximum likelihood and (B) maximum parsimony ancestral character reconstructions for the presence/absence of axillary inflorescences. Pie charts at nodes in A indicate the probability of a given state at that node—present (black) or absent (white). Letters in boxes at nodes in B indicate the state(s) reconstructed at that node—A=absent and P=present—coloration on branches indicates the state(s) reconstructed along the branch and matches the color scheme in A.

**Appendix S.16.** (A) Maximum likelihood and (B) maximum parsimony ancestral character reconstructions for flower number per raceme. Pie charts at nodes in A indicate the probability of a given state at that node—>20 (black), 10-20 (gray) or <10 (white). Letters in boxes at nodes in B indicate the state(s) reconstructed at that node—A=>20, B=10-20, and C=<10—coloration on branches indicates the state(s) reconstructed along the branch and matches the color scheme in A.

**Appendix S.17.** (A) Maximum likelihood and (B) maximum parsimony ancestral character reconstructions for flower color. Pie charts at nodes in A indicate the probability of a given state at that node—literal (white, pale yellow/green, orange, or purple). Letters in boxes at nodes in B indicate the state(s) reconstructed at that node—W=white, Y=pale yellow/green, O=orange, and P=purple—coloration on branches indicates the state(s) reconstructed along the branch and matches the color scheme in A.

**Appendix S.18.** (A) Maximum likelihood and (B) maximum parsimony ancestral character reconstructions for fruit color. Pie charts at nodes in A indicate the probability of a given state at that node—red synchronous (red), black synchronous (black), black sequential (blue), or dry fruits (brown). Letters in boxes at nodes in B indicate the state(s) reconstructed at that node—R=red synchronous, B=black synchronous, S=black sequential, D=dry—coloration on branches indicates the state(s) reconstructed along the branch and matches the color scheme in A.

**Appendix S.19.** (A) Maximum likelihood and (B) maximum parsimony ancestral character reconstructions for presence/absence of thorns. Pie charts at nodes in A indicate the probability of a given state at that node—present (black) or absent (white).). Letters in boxes at nodes in B indicate the state(s) reconstructed at that node—A=absent and P=present— coloration on branches indicates the state(s) reconstructed along the branch and matches the color scheme in A.

**Appendix S.20.** Ancestral reconstruction of thorns on various topologies for clade VI, section *Armatum*. Visualization scheme matches that of Appendix S.19.

## Footnotes

1 Manuscript received___; revision accepted___

## Notes

### Competing Interest Statement

The authors have declared no competing interest.

