## Supplementary figures and images for "Phylogeny, classification, and character evolution of tribe Citharexyleae (Verbenaceae)"

### Appendix S3

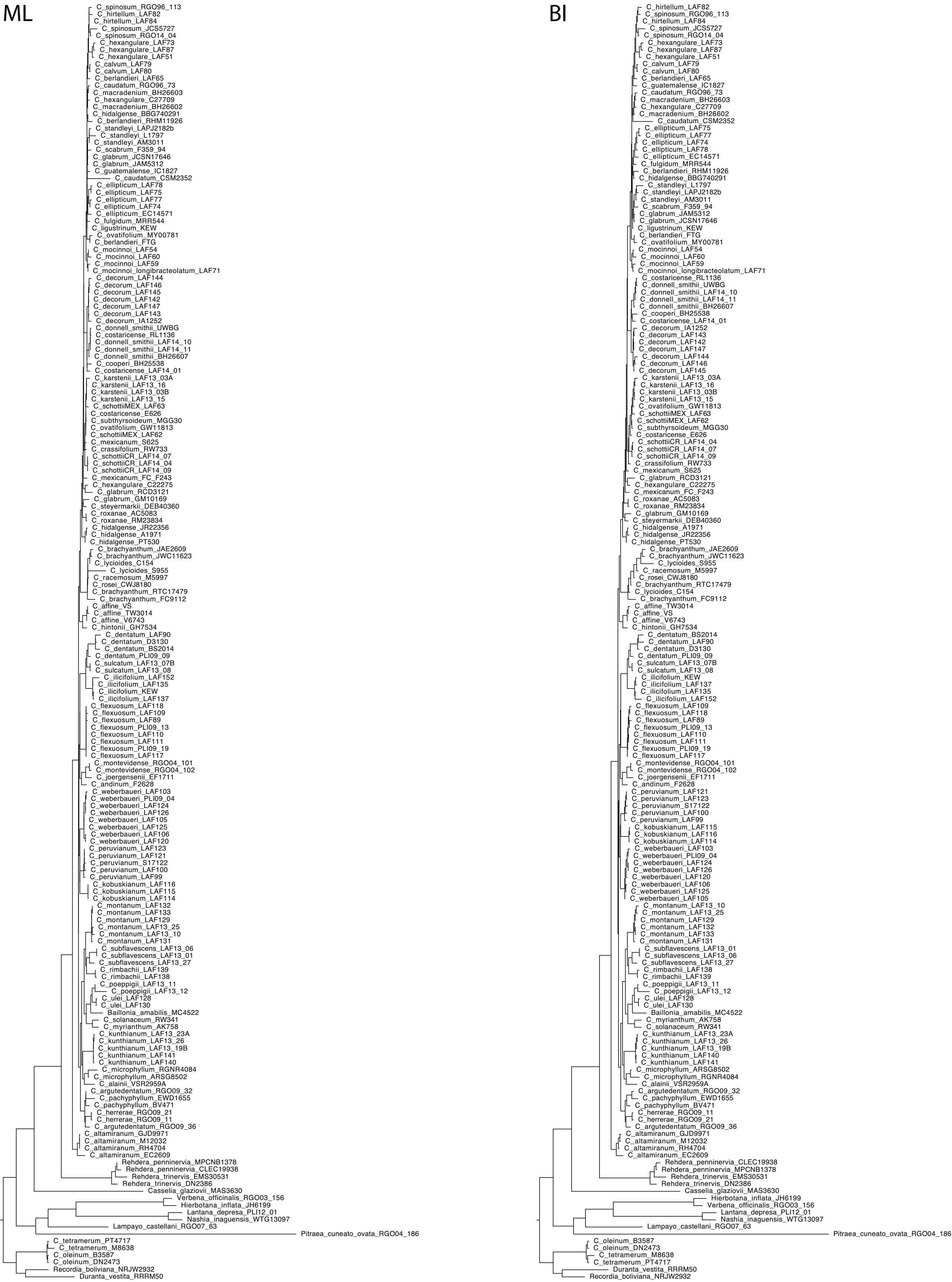

### Appendix S4

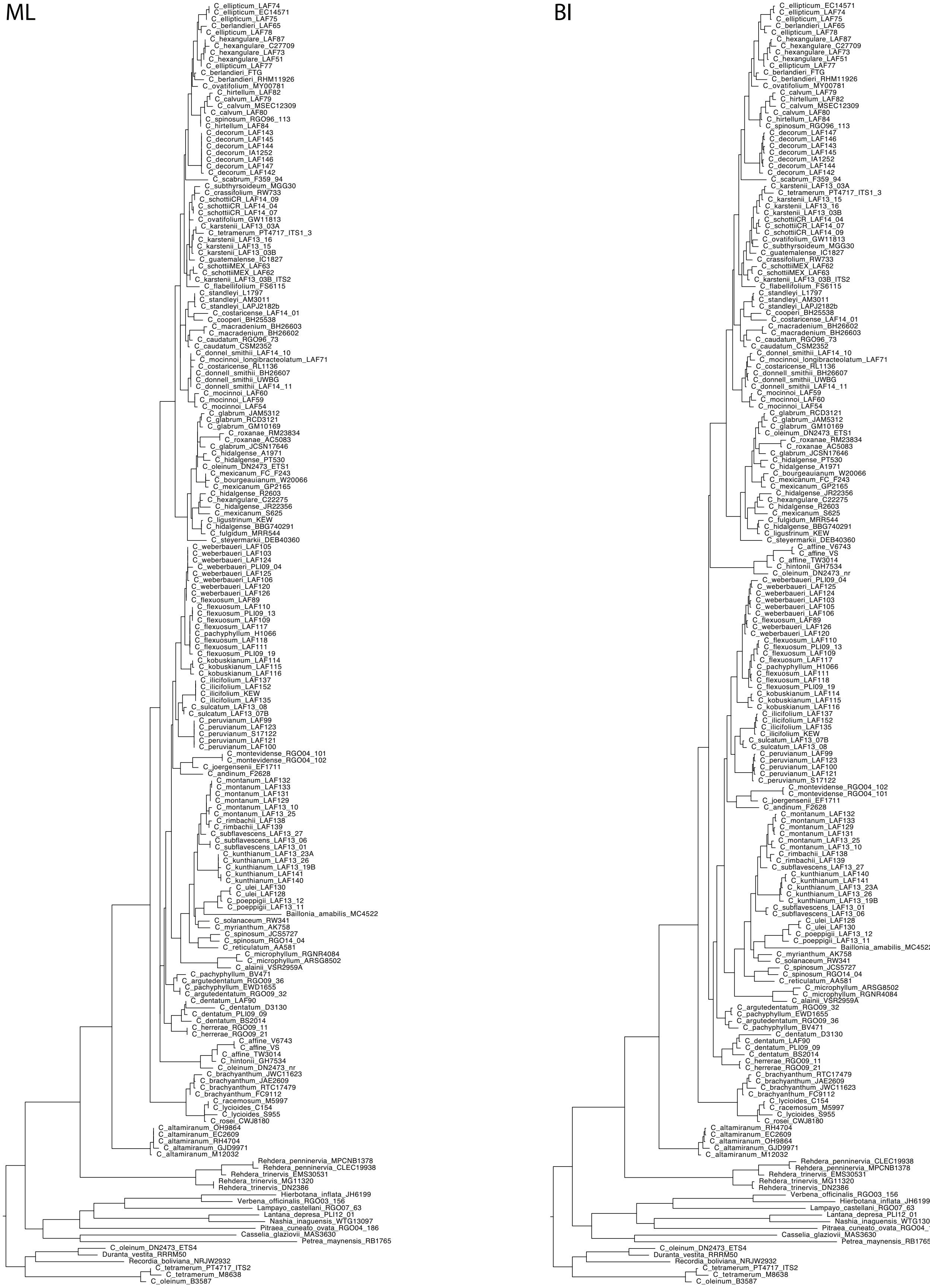

### Appendix S5

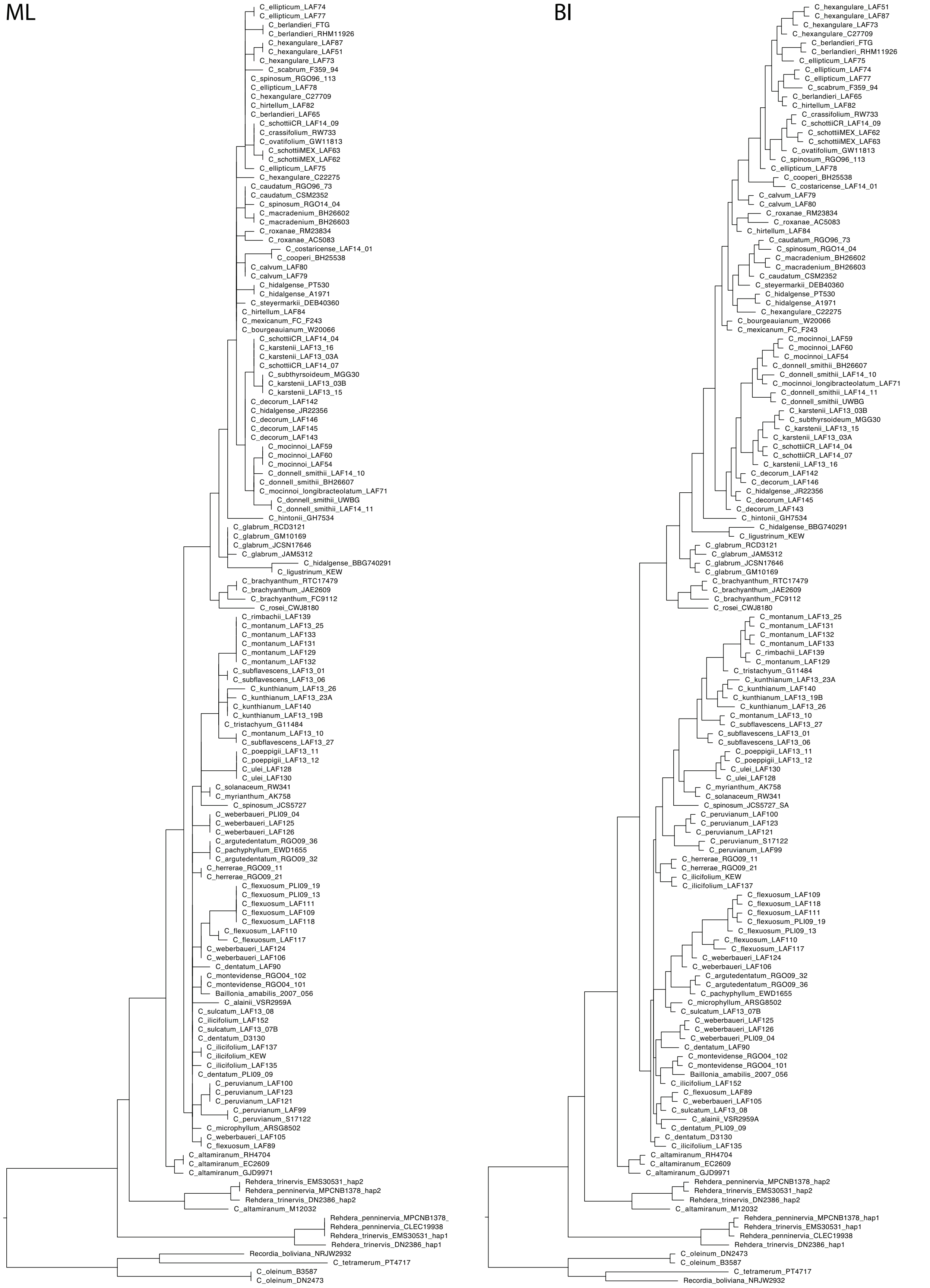

### Appendix S6

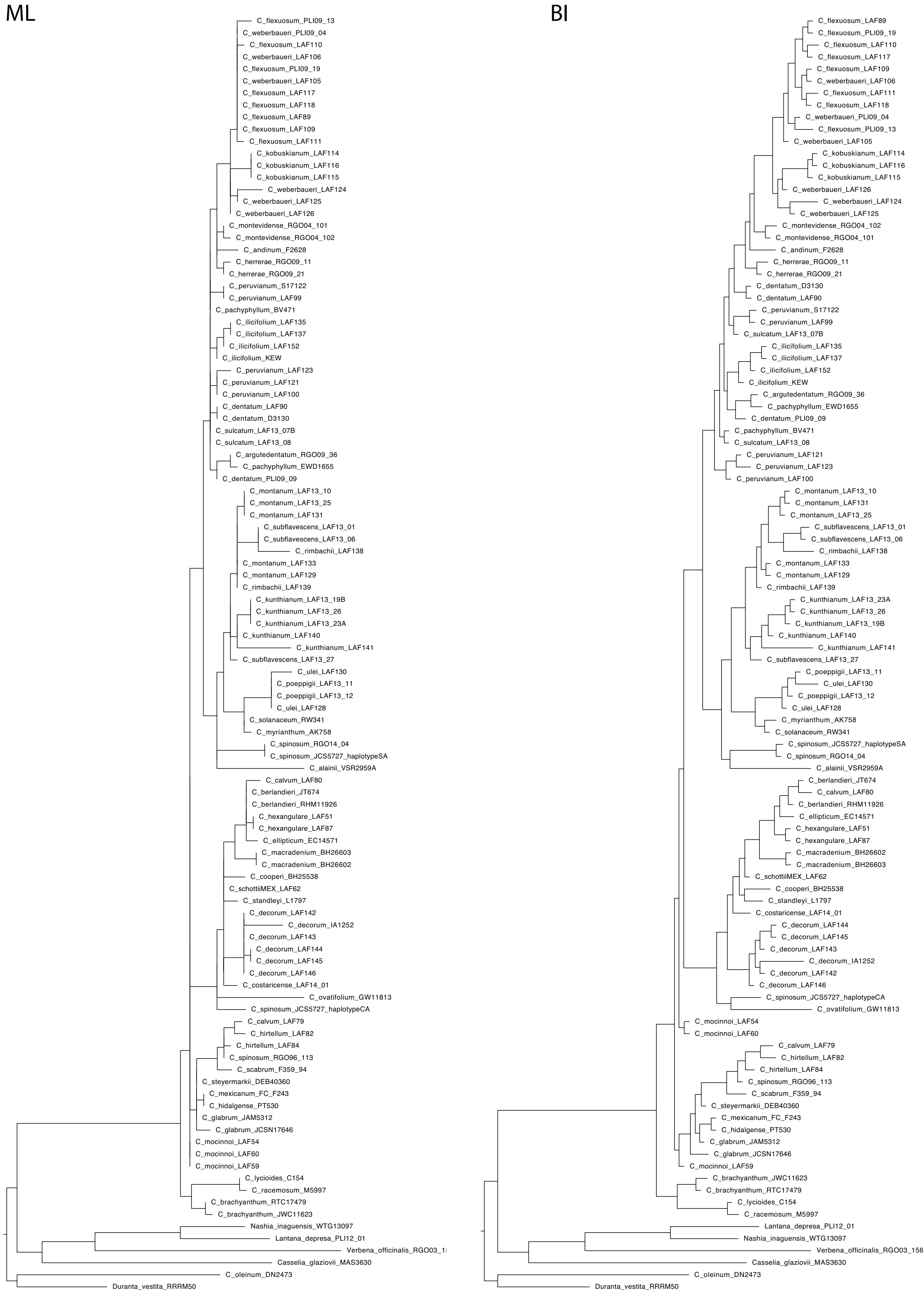

### Appendix S7

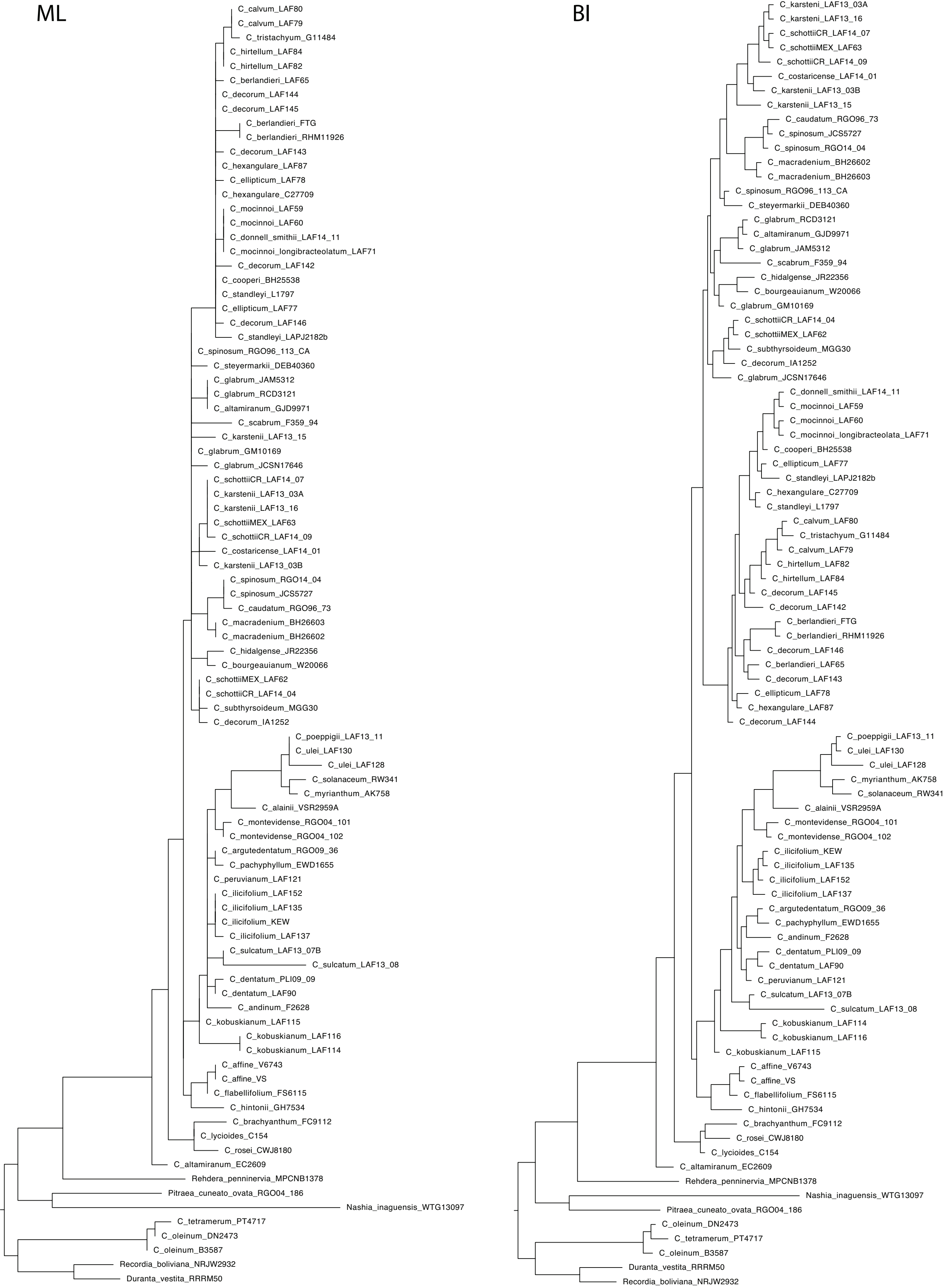

### Appendix S8

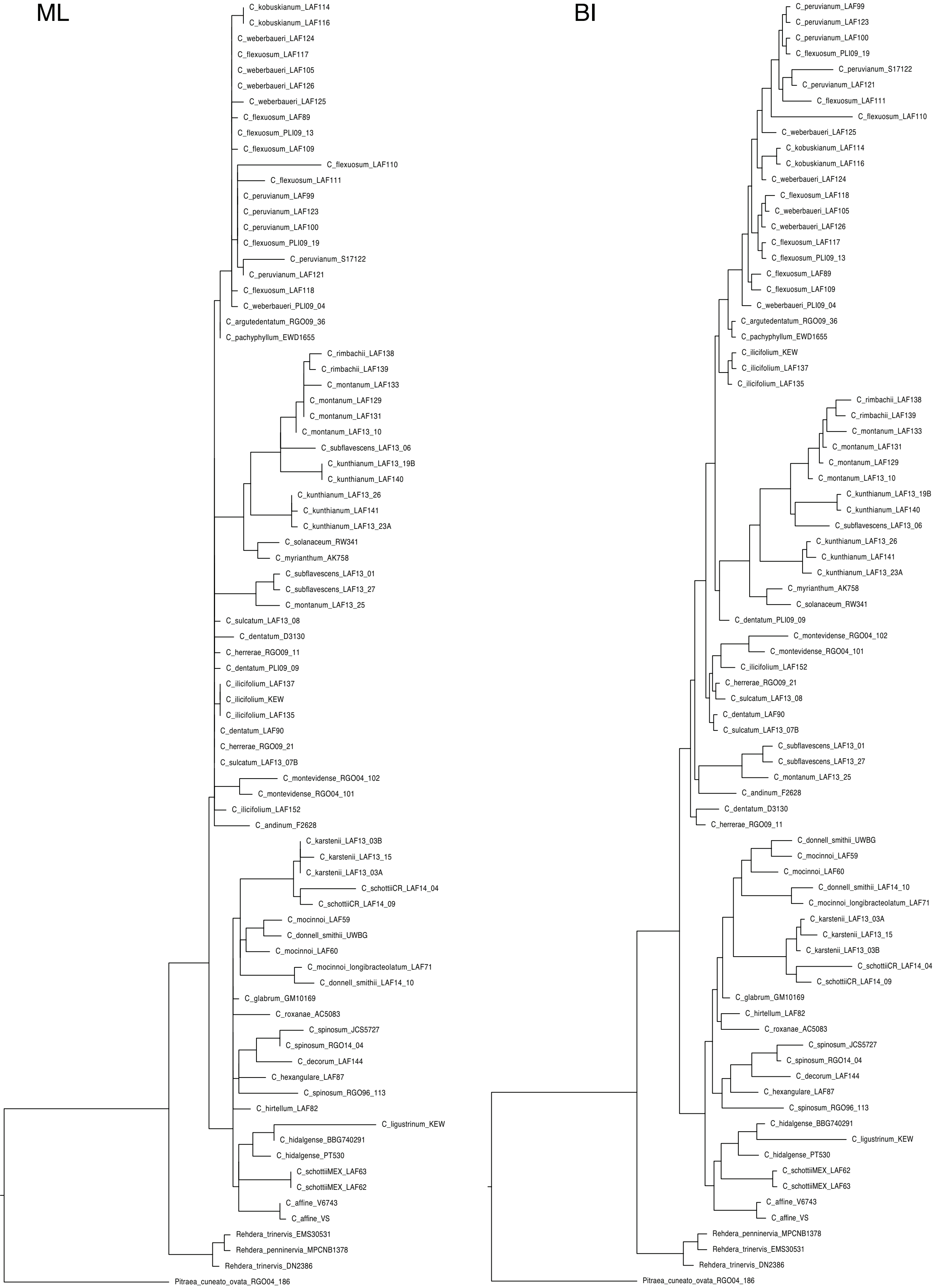

### Appendix S9

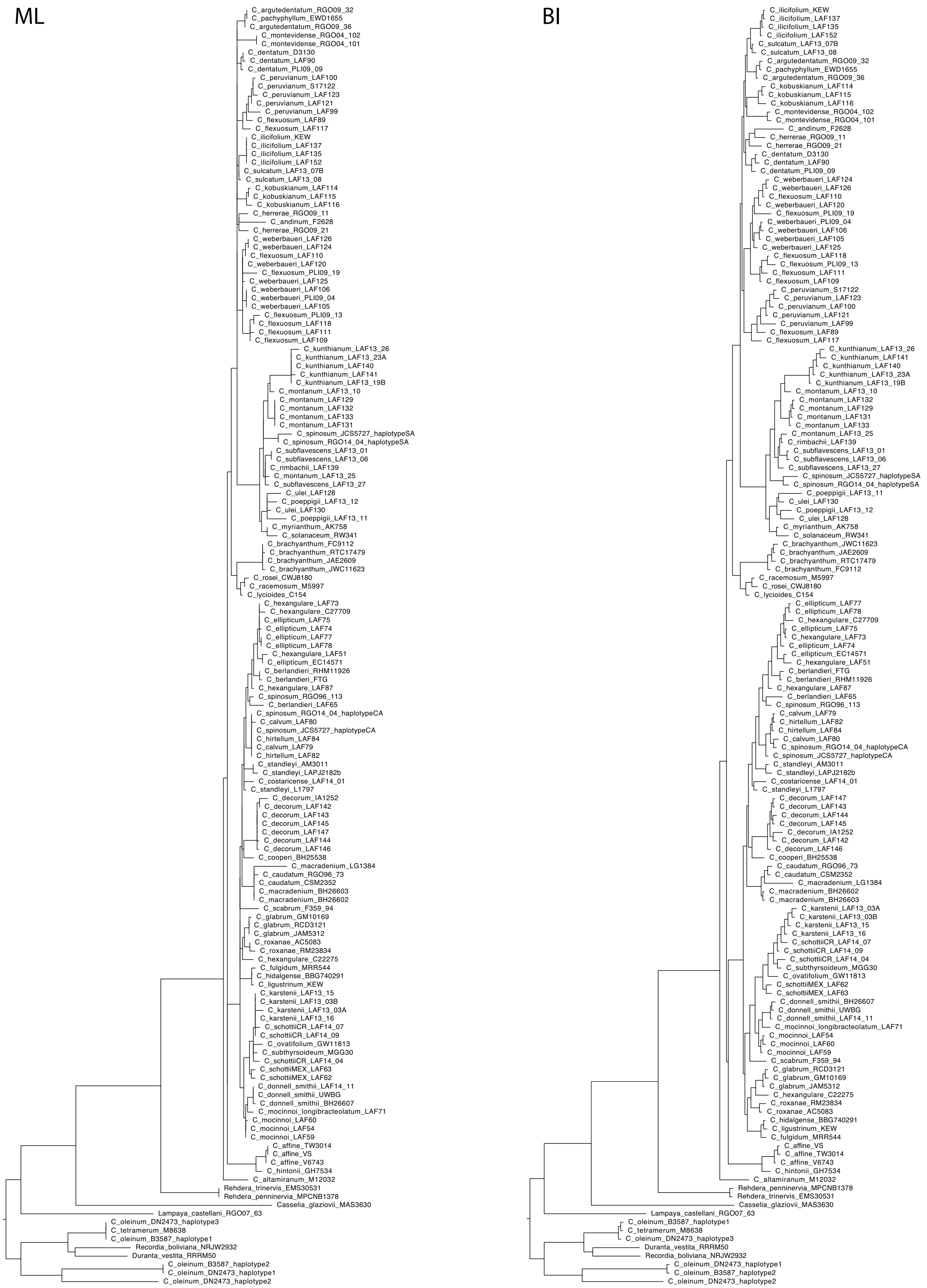

### Appendix S10

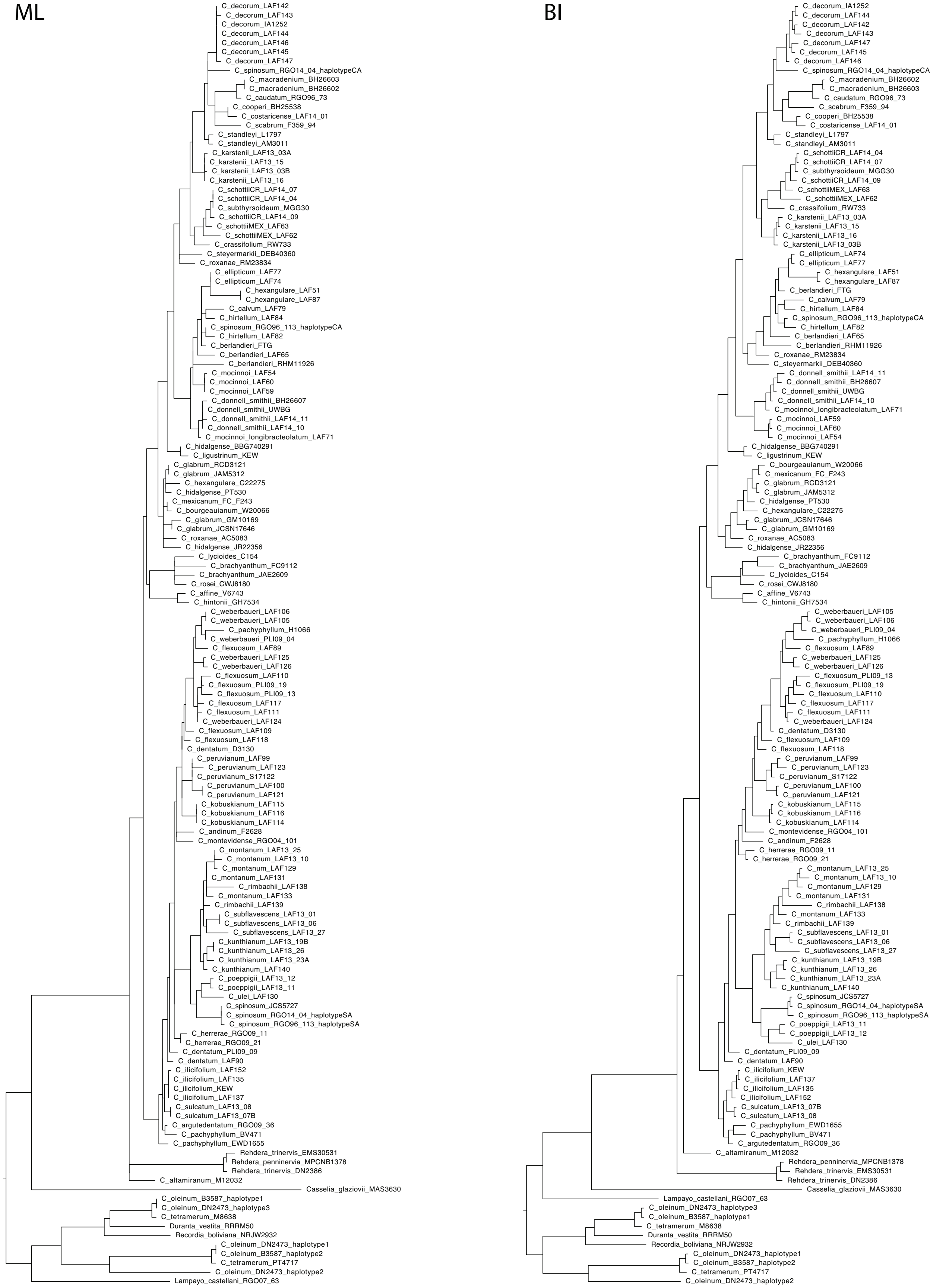

### Appendix S11

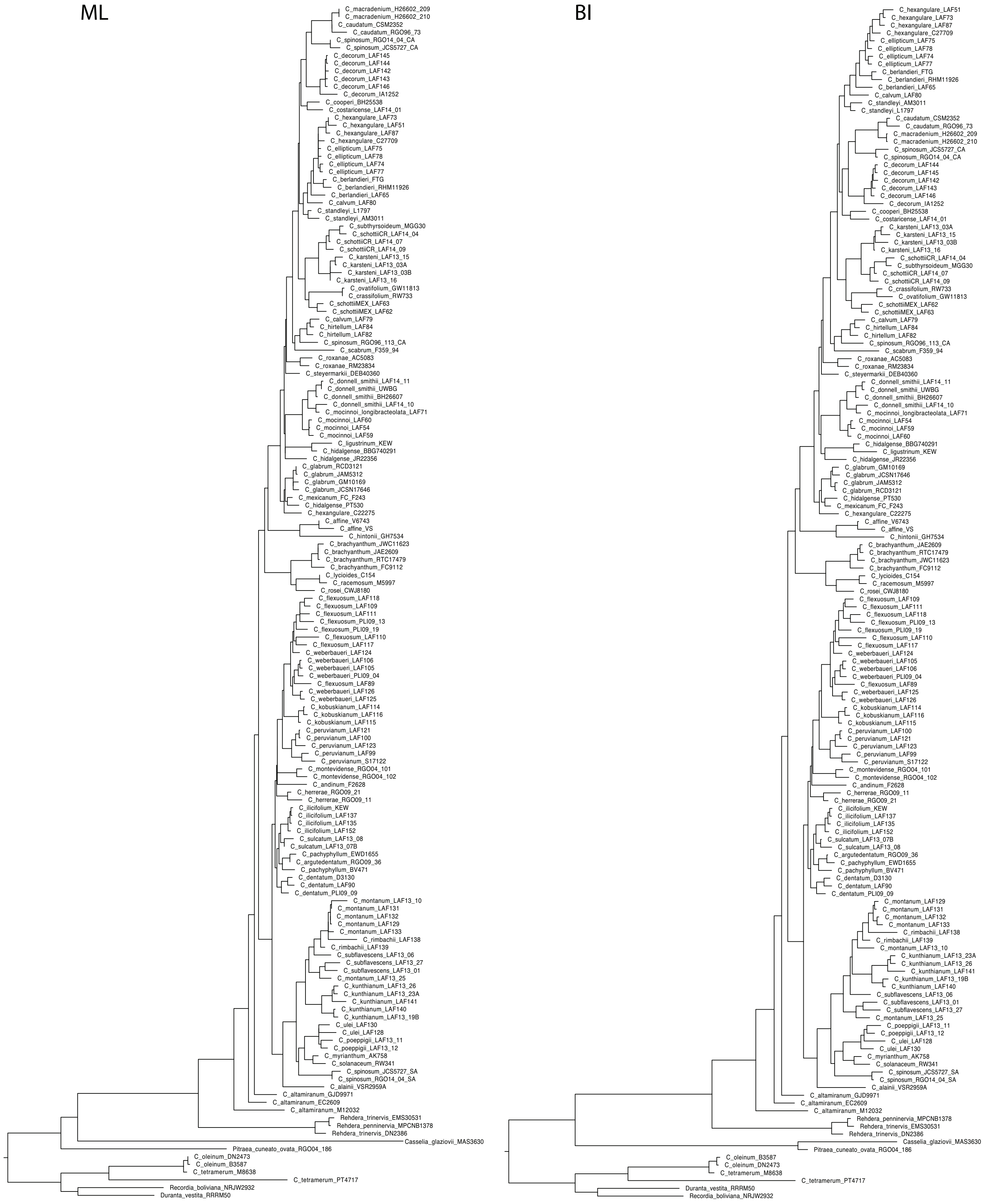

### Appendix S12

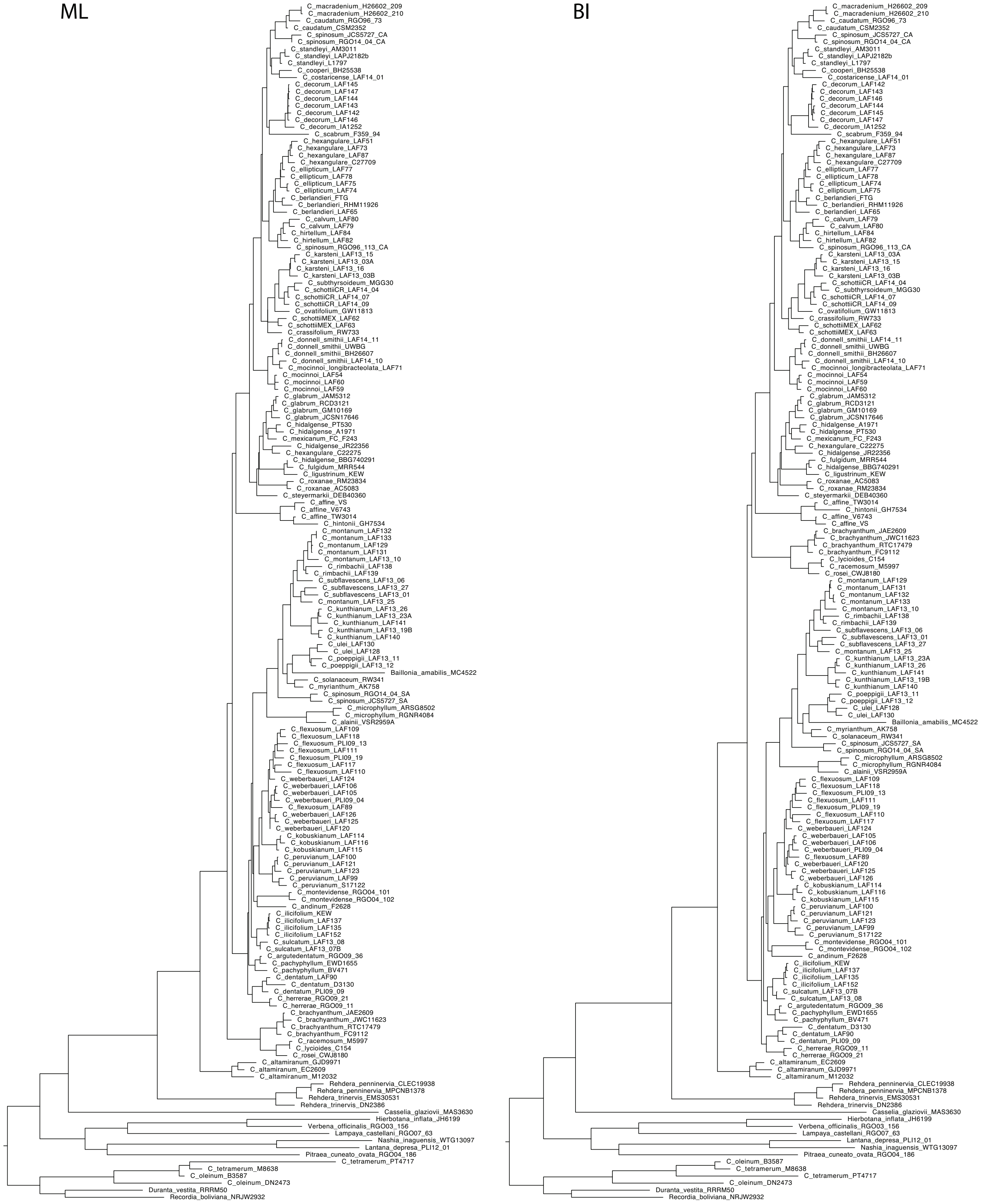

### Appendix S13

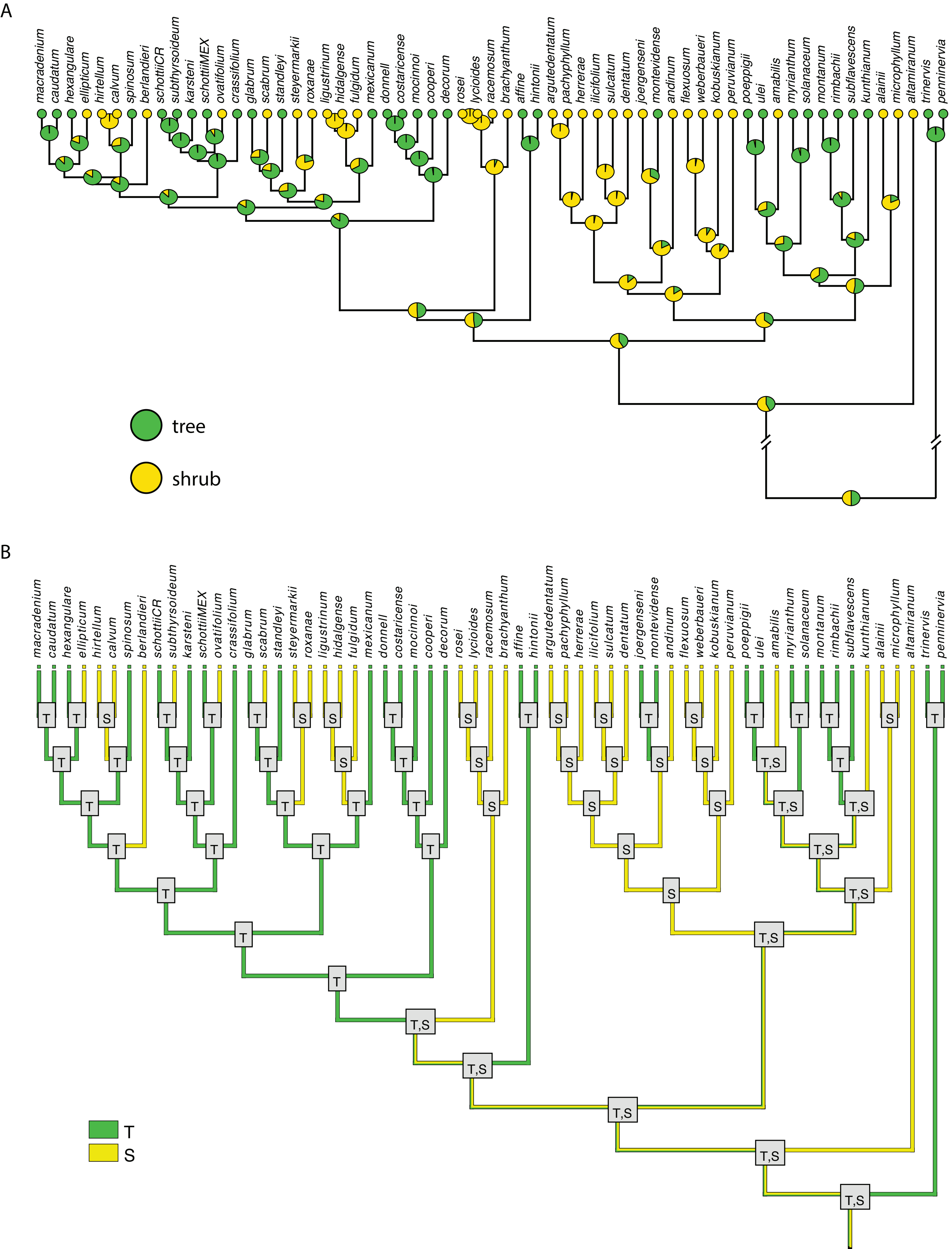

### Appendix S14

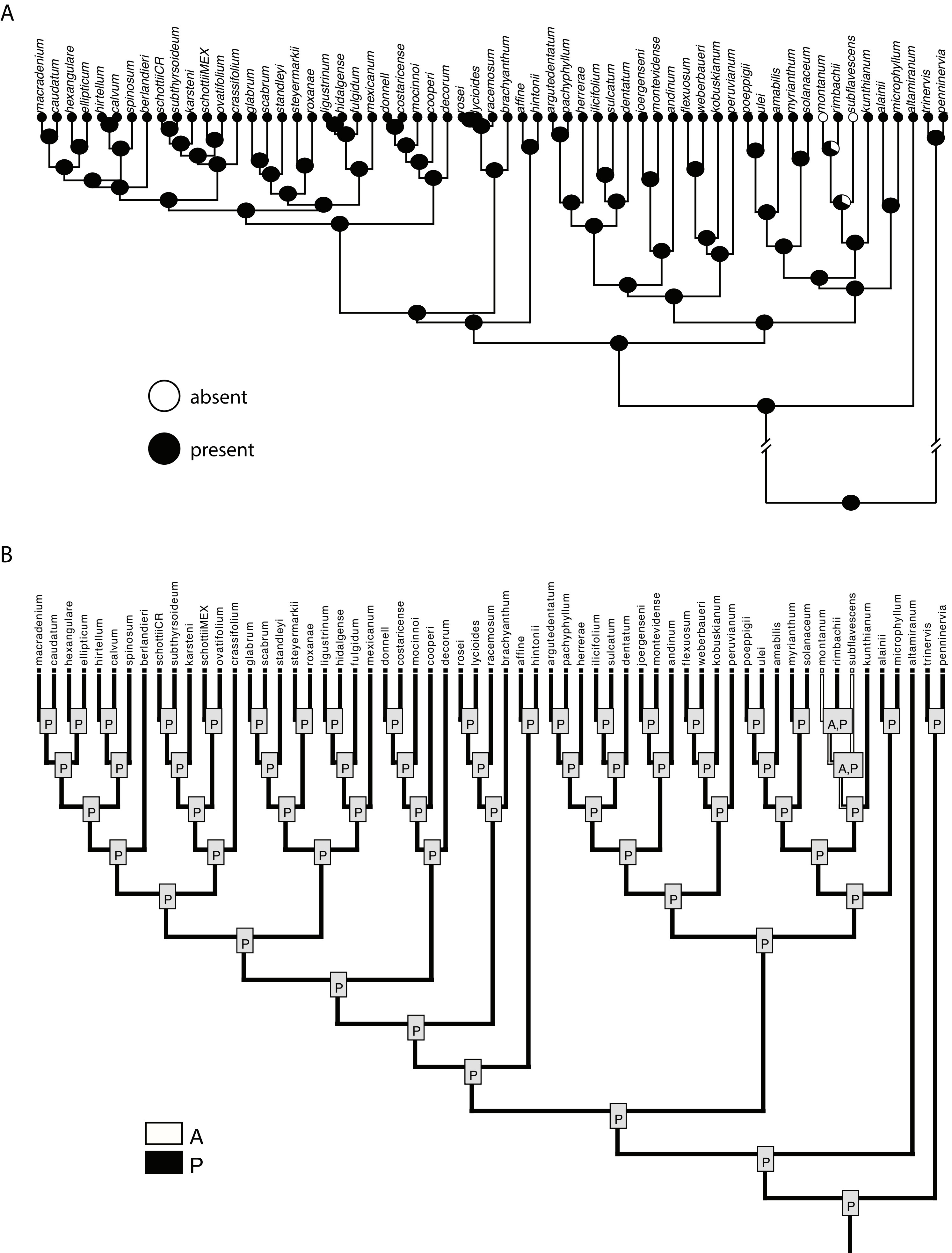

### Appendix S15

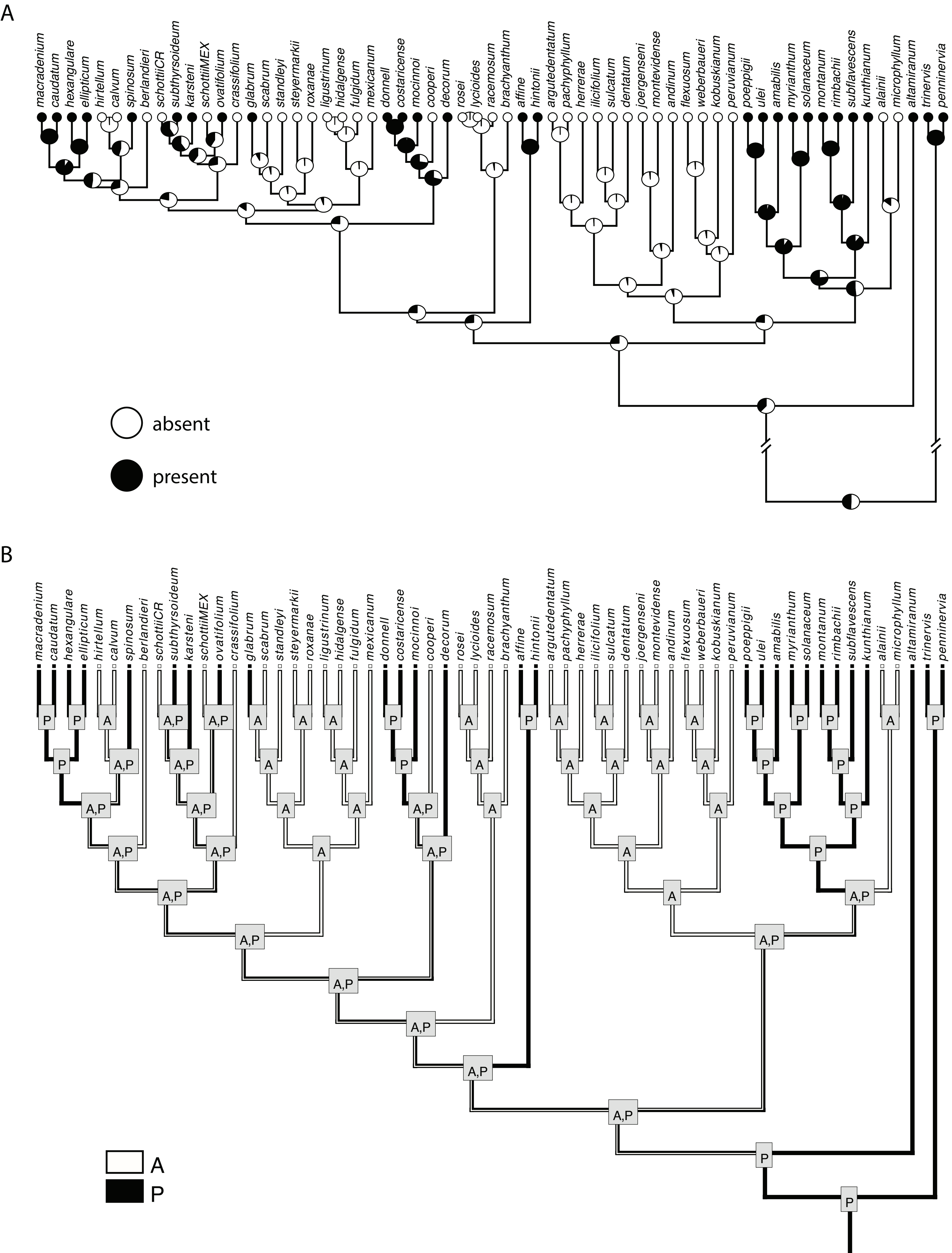

### Appendix S16

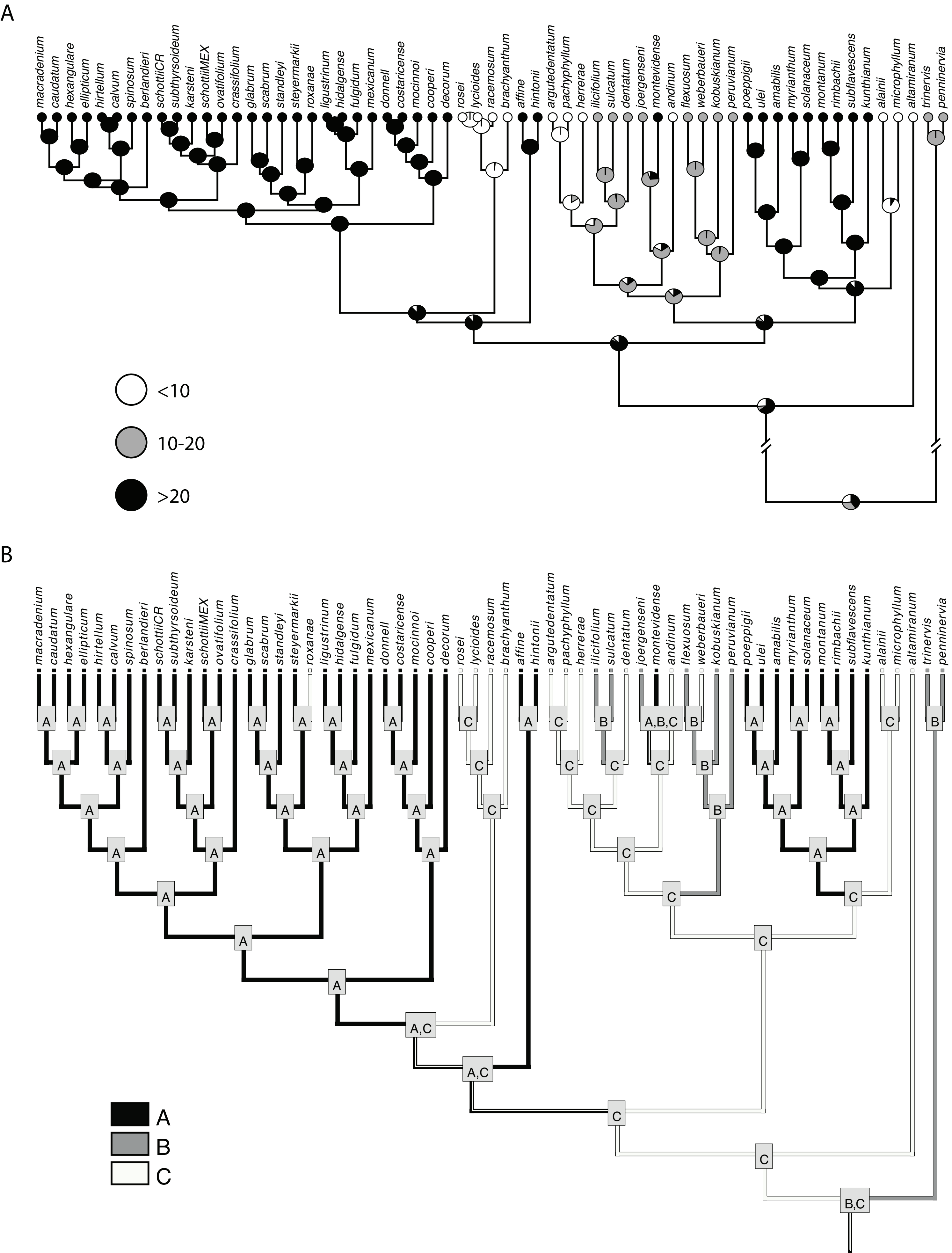

### Appendix S17

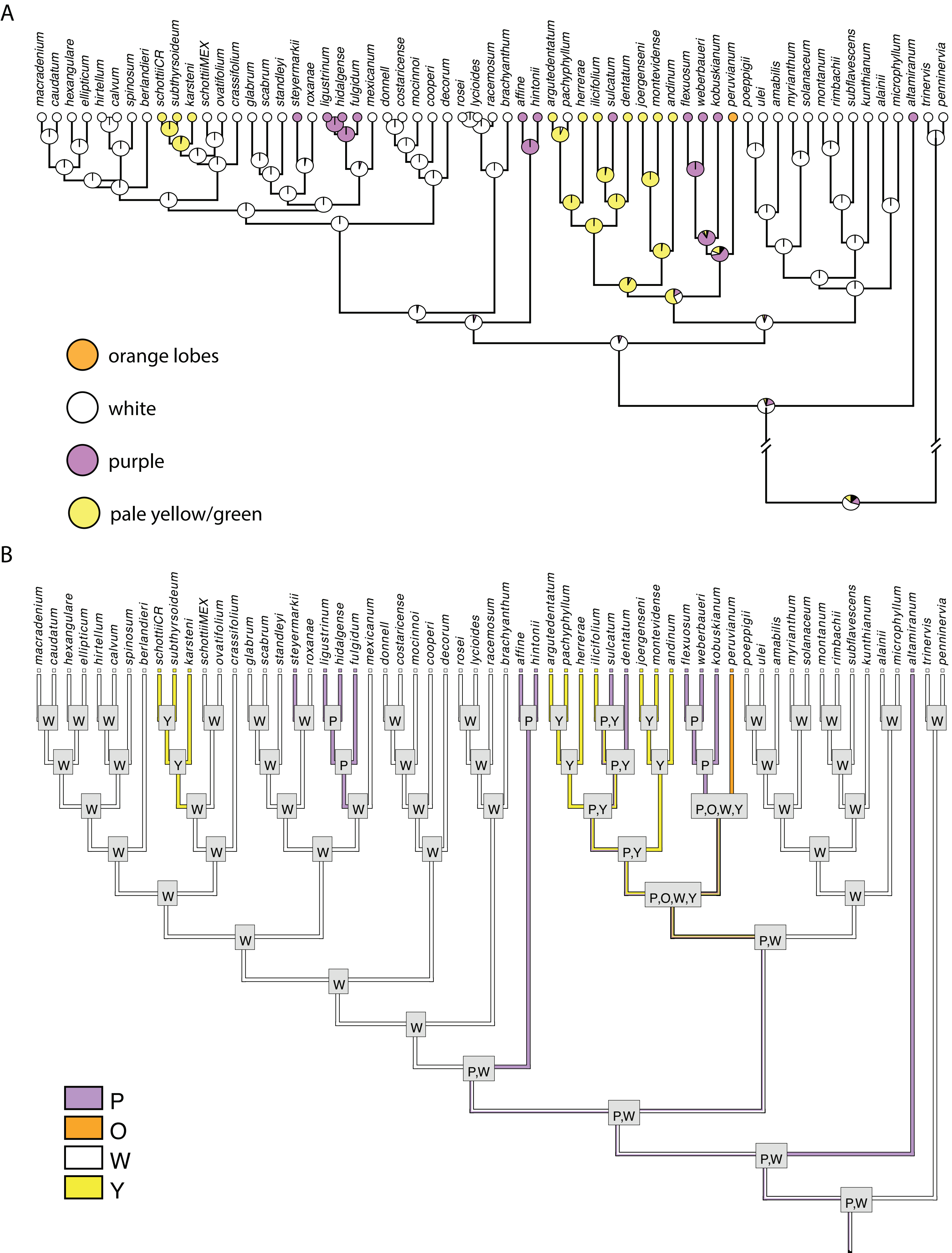

### Appendix S18

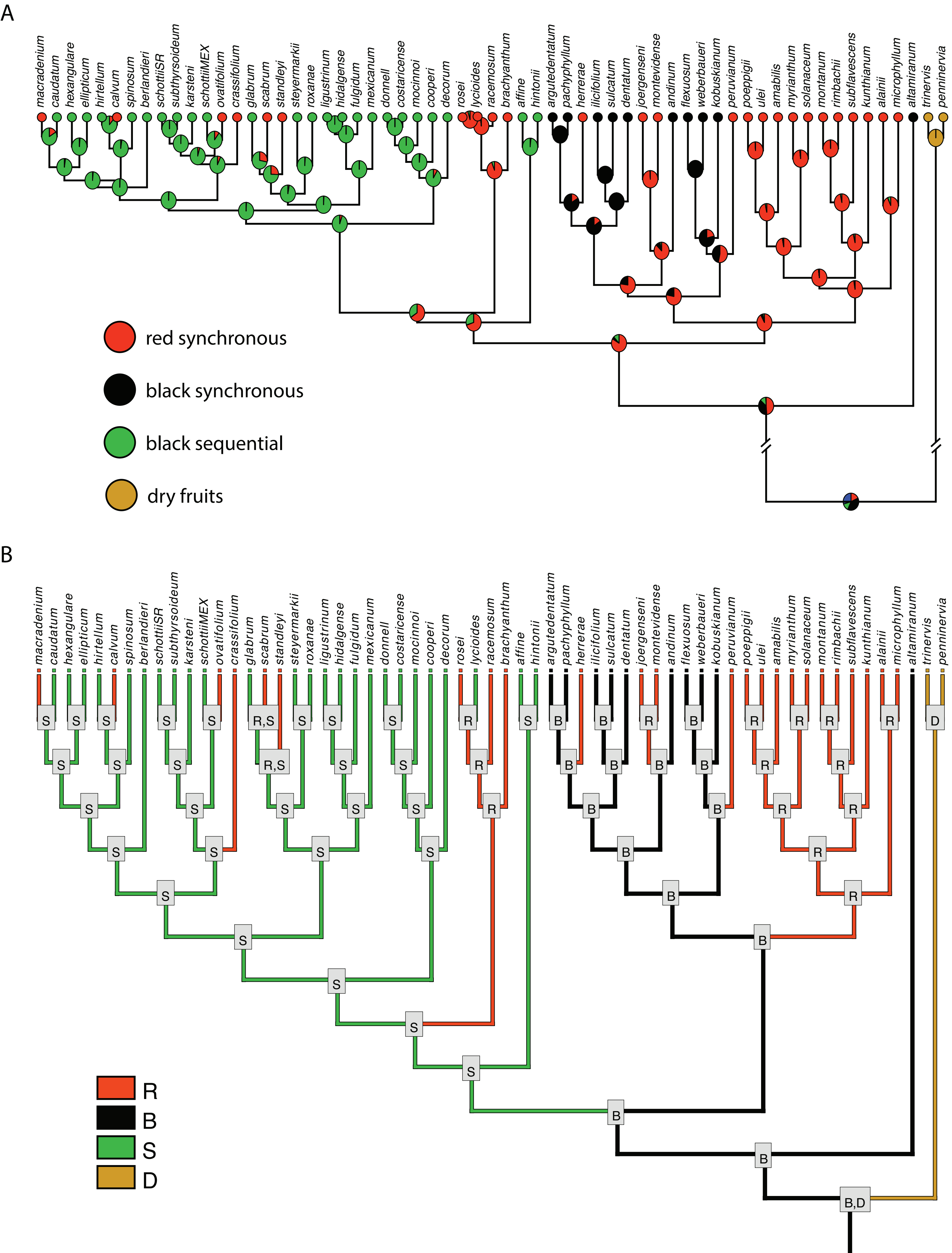

### Appendix S19

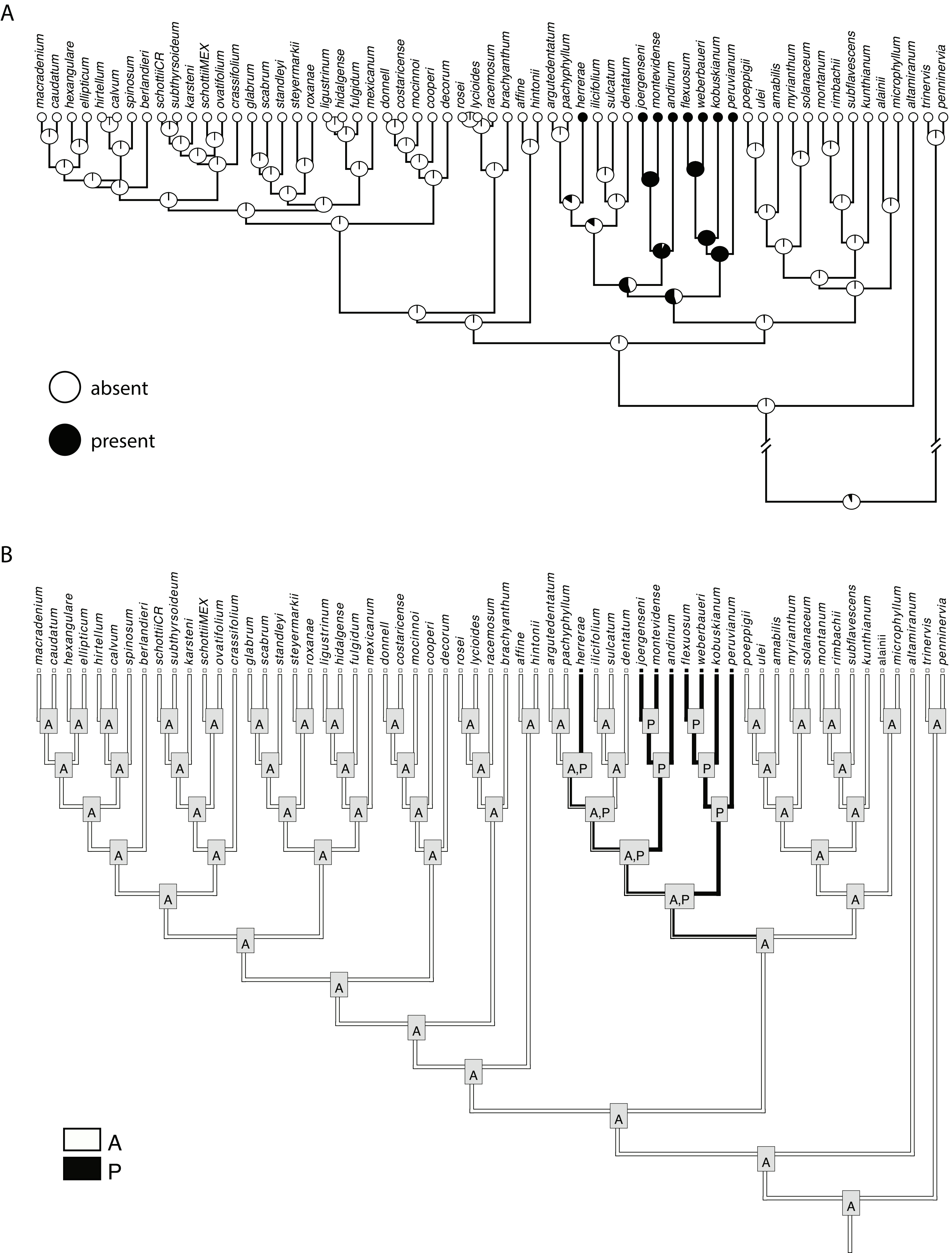

### Appendix S20

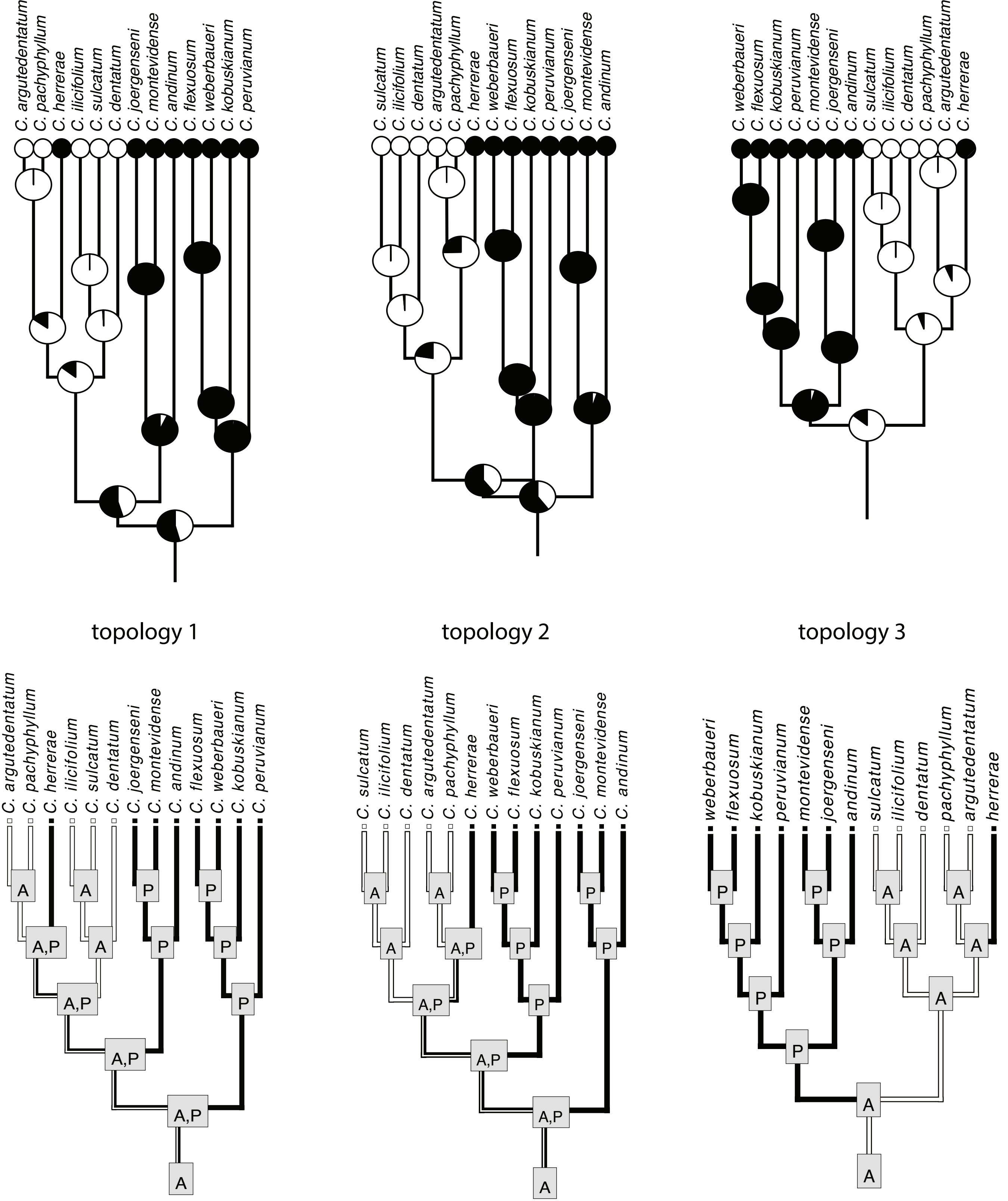
